# Multimodal imaging maps spinal glymphatic transport in health and its disruption after injury

**DOI:** 10.64898/2026.09.16.752122

**Authors:** Jaspreet Kaur, Michael J. Giannetto, Devin T. Wong, Douglas H. Kelley, Pia Weikop, Maiken Nedergaard

## Abstract

Spinal cord injury (SCI) is an incurable neurological condition in which post-traumatic edema contributes to secondary ischemic damage because the spinal cord is confined within the rigid vertebral canal. However, glymphatic transport in the healthy and injured spinal cord remains poorly characterized. We hypothesized that spinal cerebrospinal (CSF) and glymphatic transport is shaped by anatomy, administration route, and molecular size of the tracer; is disrupted after SCI; and is further impaired by loss of aquaporin-4 (AQP4).

CSF tracers were administered by cisterna magna or lumbar intrathecal injection to anesthetized adult C57BL/6 mice. Dynamic contrast-enhanced MRI, ex vivo fluorescence imaging, and histology were used to characterize tracer distribution, influx, and clearance.

Constrictions of the spinal subarachnoid space at C3-C7, T6-T12 and L5-S1 spinal segments accelerated contrast-agent dispersion, demonstrating that local anatomy regulates spinal CSF transport. Tracer influx into the cord itself occurred primarily along periarterial pathways.Tracer distribution depended on the administration route: cisterna magna injection preferentially labeled white matter in the upper spinal cord, whereas lumbar injection preferentially labeled grey matter in the lower spinal cord. Gadobutrol and ovalbumin penetrated and dispersed throughout the parenchyma, whereas fibrinogen remained confined to the meninges. Under physiological conditions, intraparenchymal tracers cleared within 24 h.

SCI was modeled using a minimally invasive NMDA-induced lesion at T11 that produced focal necrosis while preserving the dura and vertebral structures. Glymphatic transport, lesion volume, and motor recovery were compared between wild-type and AQP4-knockout mice using imaging, the Basso Mouse Scale, and voluntary wheel running. After SCI, intrathecal contrast propagated more rapidly, especially at T7-T8 segments and at the S1 segment, but tissue clearance was profoundly impaired, with tracer retention persisting for up to seven days. AQP4-knockout mice showed greater clearance impairment, larger lesions, and poorer motor recovery than wild-type controls compared at 24 h.

These findings define major determinants of spinal glymphatic transport and show that accelerated tracer propagation after SCI does not indicate effective clearance. Instead, SCI produces glymphatic dysfunction that is exacerbated by loss of AQP4 and associated with greater tissue damage and functional impairment. The results extend the glymphatic framework to the spinal cord and identify fluid-clearance pathways as potential therapeutic targets for limiting edema and secondary injury. Future studies should determine whether restoring glymphatic function improves neurological recovery after SCI.

## Introduction

Spinal cord injury (SCI) is a significant global health issue, with an estimated annual incidence of over 900,000 cases worldwide (GBD, 2016) and more than 15 million individuals living with the long-term consequences of SCI (WHO, 2024). Clinically, corticosteroids such as methylprednisolone are used as a first-line treatment following SCI (Fehlings et al, 2017). However, these drugs are associated with a range of adverse side effects and their efficacy remains controversial (Sámano et al., 2016; Liu et al, 2019). Clinical management remains largely supportive and is focused on hemodynamic stabilization, surgical decompression when indicated, and prevention of complications, followed by prolonged rehabilitation (Kaur et al, 2016). A principal driver of secondary injury is swelling of the spinal cord within the rigid bony canal, which compresses the microcirculation and reduces perfusion (Hu et al, 2023). Targeting edema may represent a critical therapeutic opportunity to limit progressive injury and preserve spinal cord function (Samano et al, 2016). Recent studies of brain ischemic edema induced by arterial occlusion or cardiac arrest have demonstrated that glymphatic transport, specifically periarterial influx of CSF, is a major contributor to acute edema formation (Mestre et al, 2020; Du et al, 2022). Moreover, cerebral edema after traumatic brain injury may stem from impaired glymphatic fluid efflux (Hussain et al, 2023). These findings suggest that edema is not merely a local event triggered by blood-brain barrier dysfunction, but that global disruption in fluid transport contributes to it (Hussain & Nedergaard, 2024).

Our understanding of glymphatic flow in the spinal cord remains limited (Wang et al, 2022), although evidence suggests an active glymphatic-like clearance mechanism operates within the spinal cord (Gonuguntla and Herz, 2023; Hou et al, 2026). Jacob and colleagues have demonstrated the CSF drainage anatomy of the spinal vertebral column using 3D imaging of decalcified iDisco^+^ (Jacob et al, 2019). Subsequently, they developed 3D imaging of brain and vertebral lymphatic vasculature and drainage, providing evidence supporting that CNS-associated lymphatic system regulates waste clearance and immune surveillance within CNS tissues (Jacob et al, 2020; 2022).. Yet, an understanding of how SCI impacts this critical fluid transport system is unknown.

Aquaporin-4 (AQP4) channels are well-known to facilitate CSF influx from perivascular spaces into the brain parenchyma (Iliff et al, 2013; Jessen et al, 2015). Previous studies have shown the role of AQP4 channels in the spinal cord (Pan et al, 2019, 2022; Kitchen et al, 2020; Berliner et al, 2023), and their role in SCI increasingly studied. Yet, no studies have directly addressed the role of AQP4 in CSF flow dynamics after the SCI (Berliner et al, 2023; Garcia et al, 2023)

To address these gaps, we asked: (1) Is spinal cord glymphatic transport polarized? Specifically, does CSF influx occur along periarterial spaces with efflux along perivenous and nerve-associated pathways, as observed in brain? (2) Does AQP4 facilitate CSF transport and tracer dispersion into the spinal cord under physiological conditions? If so, how is its distribution and function adapted to the unique anatomical and physiological context of the spinal cord? (3) How does SCI affect CSF transport in the spinal cord? Does SCI impair glymphatic flow, and how does this contribute to the disruption of fluid flow and tissue pathology? (4) Does AQP4 deletion alter CSF transport and functional recovery after SCI? If AQP4 is critical to glymphatic flow, does its absence exacerbate or mitigate post-injury fluid dynamics, and recovery?

These questions aim to unravel the role of glymphatic transport and AQP4 channels in spinal cord physiology and pathology, potentially opening new avenues for therapeutic strategies targeting fluid dynamics in SCI.

## Results

### CSF Flow Dynamics in the Subarachnoid Space Surrounding the Spinal Cord

To determine the optimal approach for studying CSF flow in the mouse spinal cord, we first compared tracer delivery via a cannula implanted in the cisterna magna (CM) or intrathecally at the lumbar spinal cord (L1 spinal level). After a 2.5-min baseline acquisition, the CSF contrast agent, gadobutrol (605 Da), was administered, and CSF dynamics were mapped in vivo using dynamic contrast-enhanced MRI (DCE-MRI) over the subsequent 60 min in ketamine (100 mg/kg) and xylazine (10 mg/kg; K/X) anesthetized mice **Fig. 1a**). Unless noted, the mice were K/X anesthesized in all experiments.

**Fig. 1.**
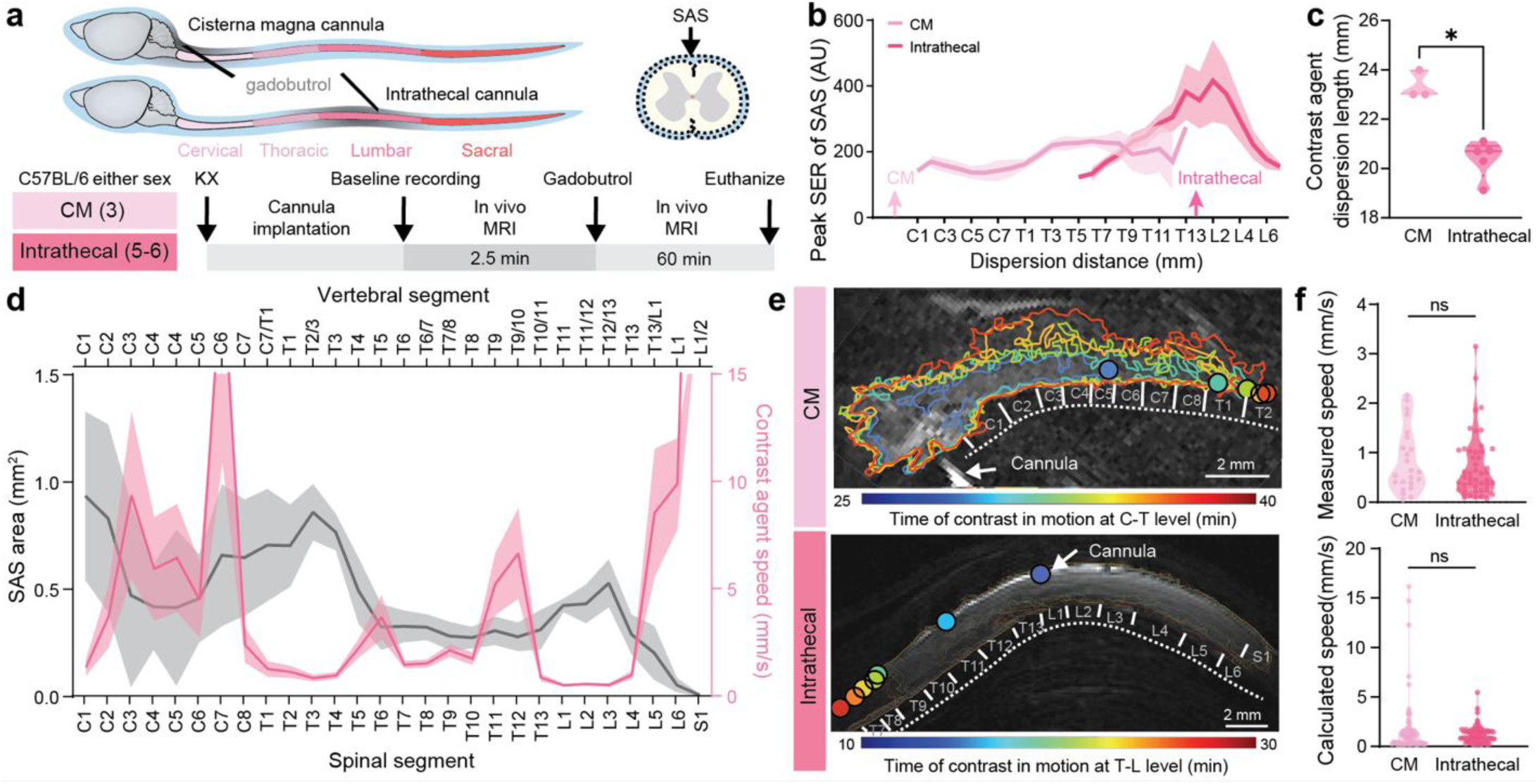
Cerebrospinal fluid flow in the spinal subarachnoid space assessed by magnetic resonance imaging. **a)** Experimental design included a 2.5 min baseline acquisition followed by dynamic contrast-enhanced MRI (DCE-MRI) in adult ketamine (K, 100 mg/kg)/Xylazine (X, 10 mg/kg) anesthetized mice of both genders. Gadobutrol (12.5 mM, 12 µl, 1.5 µl/min, 605 Da) was delivered into the cisterna magna (CM) or intrathecally at the spinal cord level (L1-L2), and images were acquired every 2.5 min throughout the 60 min imaging period. **b)** Contrast dispersion across spinal levels (C1–T13 for CM; T6–L6 for intrathecal injection) displayed as peak signal enhancement ratio (SER) during the 60 min recordings (one-way ANOVA, CM vs intrathecal, **P = 0.0063). c) Contrast agent dispersion length for both CM and intrathecal injections (Mann-Whitney test, CM vs intrathecal, P=0.035). **d)** Line graphs showing SAS area (in grey n=9) and contrast agent speed (in pink) within SAS highlighting 3 major bottlenecks between C3-C7, T6-T12 and L5-S1. **e)** Time-resolved images illustrating expansion of high-contrast regions after CM and intrathecal delivery. Edges of these regions were manually tracked to quantify contrast propagation speed. Circles mark locations where speeds came from direct measurements. Scale bar = 2 mm. **f)** Measured tracer displacement rates in the SAS showed no significant difference between CM and intrathecal delivery (two-sample t-test, P = 0.535). Calculated flow speeds across spinal segments likewise did not differ significantly between routes (two-sample t-test, P = 0.102, n= 3-6).

We first assessed the CM and intrathecal delivery routes by analyzing contrast arrival and dispersion along the spinal subarachnoid space (SAS) which was quantified as a signal enhancement ratio (SER) (**Fig. 1 and Supplementary Fig. 1**). Following CM injection, the contrast agent rapidly entered the spinal SAS at the C1 level within 3 min and reached the T13 segment within 12 min, demonstrating efficient longitudinal transport along the spinal cord (**Supplementary Fig. 1b).** Following intrathecal injection at L1 (T11 vertebral level), tracer entered the spinal SAS bilaterally and spread longitudinally. The tracer reached the T13 and L2 segments within 6.0 ± 1.2 min and 5.4 ± 1.0 min, respectively, propagated rostrally to the T5 segment within 9 min, and spread caudally to the S1 segment within 6 min. (**Supplementary Fig. 1b)**. Overall, the contrast agent broadly dispersed into the cervico-thoracic regions after CM injection, whereas intrathecal injection distributed it along the thoraco-lumbar segments (**Fig. 1b and Supplementary Fig. 1a, b**). Notably, arrival times measured 1 mm from the injection site did not differ significantly between the routes (**Supplementary Fig. 1**), indicating comparable local contrast distribution. Yet, the two administration routes produced markedly different longitudinal distributions: contrast injected into the CM spread over a greater length of 23.33 ± 0.33 mm along the spinal cord than contrast delivered by lumbar intrathecal injection, which dispersed about 20.39 ± 0.32 mm (**Fig. 1c, Supplementary Fig. 1**). However, shorter dispersion of contrast longitudinally after intrathecal injection might be due to technical limitations, as we used a small surface coil of 23 mm for signal detection. Thus, regardless of whether the contrast agent was injected rostrally or caudally, it was cleared from the spinal CSF before it could fill the entire SAS or reach the opposite end of the spinal canal, indicating rapid local CSF efflux and turnover. The contrast injection pump was on for 8 min, and 12 µl of contrast agent was injected at a rate of 1.5 µl/min. CSF flow within the spinal SAS is not uniform along the neuraxis and is shaped by the local geometry of the central canal, cardiac and respiration-driven pulsatility, nerve roots and denticulate ligaments (Yiallourou et al., 2012; Sánchez et al., 2018). While pulsatile CSF velocity has been characterized extensively in humans and large animals, and net CSF/tracer transport has been modelled in vitro and computationally (Ayansiji et al., 2023), contrast agent propagation speed in the spinal SAS of rodents at the segmental level is lacking. To address this gap and to determine how regional SAS geometry is related to local fluid transport, we quantified both the SAS area and the propagation speed of contrast agent along the spinal axis using DCE-MRI. The SAS area was minimal between C3-C7, T6-T12 and L5-S1 segments, highlighting the major bottleneck regions and demonstrating that the SAS is highly heterogeneous along the mouse spinal axis, creating regional constrictions and expansions that are likely to influence CSF flow dynamics and tracer propagation (**Fig. 1d, grey line graph**). We next quantified contrast propagation speed within the SAS relative to the spinal segment based on the DCE-MRI signal (**Fig. 1d, pink line graph**). Contrast propagation was slow at the C1 cervical region, which began to accelerate at C2 and increased between C3-C7 to 9.26±3.9 mm/s, highlighting the beginning of the first regional bottleneck, which again increased between T6-T12 to 4.8±1.13 mm/s, corresponding to a decreased SAS area at T6-T12. Note that even though the speed was reduced between T7-T10 segments, the overall speed between T6-T12, including T7-T10 (1.9±0.3 mm/s) segments, was higher than the minimum average speed of 0.49±0.06 mm/s. The contrast propagation was minimal between T13-L4, with a corresponding increase in area, which reached maximal levels between L5-S1 at 20.42±7.67 mm/s, with area reaching a minimum. The increased speed at C3-C7, T6-T12 and again at L5-S1 coincided with a local constriction of SAS area and emphasised the presence of major bottlenecks. Representative images in **Fig. 1e** show time-resolved contour lines of the advancing contrast front after CM or intrathecal injection. At each time point, peak signal positions along the spinal axis were identified from the contour maxima, and propagation speed was calculated from the pixel displacement between successive frames, accounting for spatiotemporal resolution (**Supplementary Fig. 2 displays the velocity of the contrast agent dispersion in each mouse**). Comparison of measured tracer displacement rates with calculated volumetric flow estimates (**Fig. 1f**; see Methods) revealed no significant difference in overall propagation speed between CM and intrathecal delivery. Thus, while contrast agent speed along the spinal cord was comparable between routes, the resulting spatial distribution of contrast depended strongly on the site of CSF delivery.

In summary, regional analysis of SAS area revealed important bottlenecks at various spinal levels where local constriction of SAS increased contrast propagation speed – identifying these regions as the critical hallmarks for fluid dynamics in the spinal cord (**Fig. 1d-f**).

### Fluid Transport within the Spinal Cord Parenchyma

Although several studies have demonstrated that CSF tracers enter the spinal cord parenchyma (Lam et al, 2017; Wei et al, 2017; Liu et al, 2022; Melin et al, 2023), systematic mapping of their longitudinal distribution and differential penetration into grey and white matter, particularly across administration routes, remains limited. To address this gap, we performed additional analyses of the in vivo MRI datasets focused on the cord itself (**Fig. 2a-f**) and complemented these with ex vivo fluorescent imaging in a separate cohort of anesthetized mice (**Fig. 2g-j**).

**Fig. 2.**
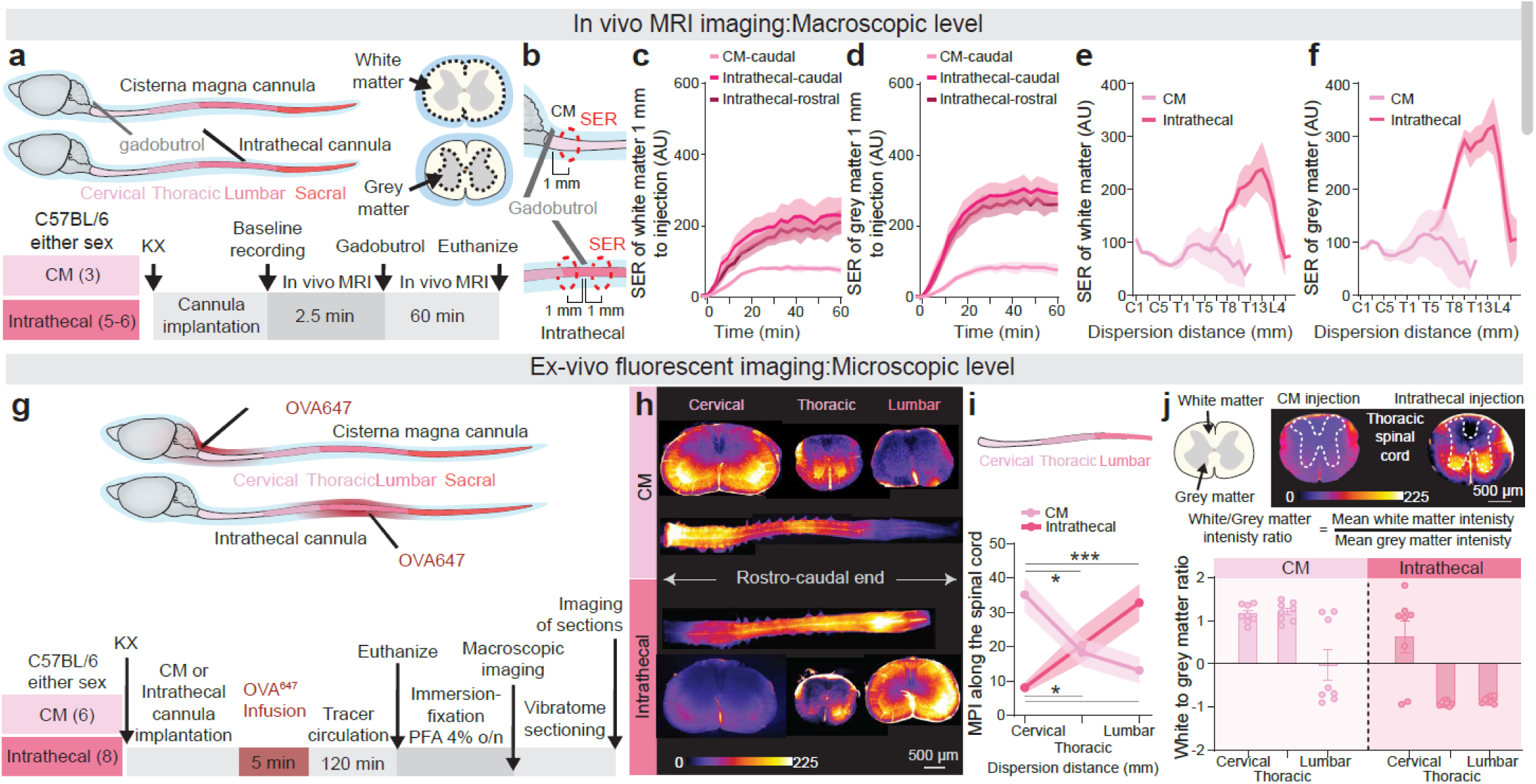
Cerebrospinal fluid tracer distribution in the spinal cord parenchyma after CM or intrathecal injection at macroscopic and microscopic levels. **a)** Baseline acquisition of 2.5 min and subsequent gadobutrol (12.5 mM, 12 µl, 1.5 µl/min, 605 Da) injection and dynamic contrast-enhanced MRI (DCE-MRI) recording for the next 60 min. **b)** Sketch of measurement of Signal enhancement ratio (SER) 1 mm from the gadobutrol injection site when delivered via CM or intrathecally. **c, d)** SER in white (c) and grey matter (d) measured 1 mm from the gadobutrol injection site when administered via CM (caudally) or intrathecal (rostrocaudally) route at 60 min (white and grey matter: Two-Way ANOVA, n=3-6, ****P<0.0001, CM vs intrathecal, n=3-5). **e, f)** After CM and intrathecal infusion, peak SER in white (d, CM vs intrathecal: ****P=0.0001) and grey matter (e, CM vs intrathecal: P=0.0001; n=3-5) across 21 (C1-T13) and 16 spinal levels (T5-S1) respectively. **g)** Schematic representation of experimental design. A fluorescent tracer, ovalbumin conjugated with Alexa Fluor 647 (OVA^647^, 10 µl, 2 µl/min, 45 KDa) was administered via the CM or intrathecally at the L1 level. Spinal cords were harvested 120 min post-injection, fixed, and processed for imaging. **h)** Macroscopic imaging (4X) of whole spinal cords and cross-sections after CM or intrathecal injection. **i)** Quantitative line-plots of dispersion distance with respect to mean pixel intensity, MPI (CM injection: n = 6, t-test, cervical vs. thoracic: P = 0.0275, cervical vs. lumbar: P = 0.0061; intrathecal injection: n = 8; t-test, cervical vs. thoracic: P = 0.037, cervical vs. lumbar: P = 0.006). **j)** Example sketch of grey and white matter area with representative sections of thoracic spinal cord with tracer distribution after CM or intrathecal injection and the formula for white to grey matter intensity ratio calculation plotted as a bar graph. Note that n refers to the number of mice used.

Following CM injection, there was a slow, gradual distribution of contrast, with lower levels of saturated peak signal enhancement ratio (SER) in both white and grey matter (**Fig. 2b-d**). Here the contrast entered the white matter at C1 segment already within 3 min (**Supplementary Fig. 1c**), with SER increasing across the C1 white matter (1 mm caudal from the injection, **Fig. 2b upper panel sketch**) over 15 min (**Fig. 2c**), whereas in grey matter its arrival was delayed, appearing at 6 min (**Supplementary Fig. 1d**) and peak SER at 17.5 min (**Fig. 2d**). In contrast, intrathecal administration produced rapid and extensive contrast distribution in both white and grey matter (**Fig. 2b lower panel sketch, c-d**). However, for intrathecal injection, contrast arrived at the white matter T13 level rostrally (1 mm from L1 injection site) within 6.0 ± 1.3 min and the L2 level caudally within 4.8 ± 0.66 min (**Supplementary Fig. 1c**). Entry into the grey matter occurred at comparable times, with contrast arriving at T13 rostrally within 6.6 ± 1.1 min and at L2 caudally within 6.6 ± 1.0 min (**Supplementary Fig. 1d**). Peak SER occurred earlier in white (12.5 min) than in grey matter (17.5 min) (**Fig. 2c, d**). Quantitative analysis across 16 spinal levels revealed distinct distribution patterns following the two injection routes. After CM injection, the contrast agent was distributed relatively uniformly throughout the spinal cord in both white and grey matter (**Fig. 2e, f**). In contrast, spinal intrathecal injection produced higher peak SER values and a bell-shaped rostrocaudal distribution: SER was highest at the injection site and gradually declined with increasing distance, resulting in the strongest enhancement in the thoraco-lumbar regions (**Fig. 2e, f**). Furthermore, the delay between contrast arrival in the SAS and its entry into parenchymal compartments was shorter after intrathecal injection, whereas CM delivery showed a longer lag, especially in grey matter (**Supplementary Fig. 1e, f**). Together, these results may suggest that intrathecal delivery was associated with greater parenchymal contrast dispersion than CM injection, particularly in the lower spinal cord.

To map CSF tracer entry across spinal levels in greater detail, we injected a fluorescent tracer at CM or at the lumbar spinal cord of anesthetized mice. After 120 min of tracer circulation, spinal cords were harvested and immersion-fixed. This approach is superior with regard to distinguishing tracer entry in grey vs white matter (**Fig. 2g**). Representative whole-mount, cross-sectional images (**Fig. 2h**) and MPI line plots (**Fig. 2i**) together showed rostro–caudal tracer dispersion from the injection site, with distinct regional patterns depending on delivery route. CM injection resulted in the highest accumulation of a tracer in the cervical region, followed by thoracic, with minimal signal in the lumbar cord. In contrast, intrathecal delivery produced maximal tracer accumulation in the lumbar, followed by thoracic and then cervical segments. Together, these findings confirmed that CSF tracer entry into spinal parenchyma follows a pronounced rostro–caudal gradient that is determined by the route of CSF delivery.

To further assess the tracer’s spread from the white into grey matter across different spinal levels and delivery routes, we quantified the regional distribution as a white-to-grey matter ratio, as shown in the example sketch of the thoracic spinal cord with formula (**Fig. 2j).** A ratio above 1 indicated predominant tracer localization in white matter; a ratio of 1 reflected equal distribution; and a ratio below 1 indicated greater accumulation in grey matter. Following CM injection, the ratio exceeded 1 in cervical and thoracic regions but was below 1 in the lumbar segment, indicating preferential distribution to white matter in the upper spinal cord. Conversely, after intrathecal injection, the ratio was below 1 in both thoracic and lumbar regions, consistent with greater tracer accumulation in grey matter. These ex vivo fluorescent imaging data (**Fig. 2j**) are consistent with the in vivo MRI data (**Fig. 2e, f**) where, for example, the SER of white matter and grey matter in the thoracic region were almost similar values for CM injection, whereas in the intrathecal injection, the SER value of grey matter was higher than the white matter in the thoracic region. These results demonstrate that tracer localization was strongly influenced by delivery route: CM injection favored tracer dispersion within white matter and localization to cervical and thoracic levels, while intrathecal injection promoted tracer diffusion into grey matter, particularly at thoracic and lumbar levels.

In summary, ex vivo spinal cord analysis **(Fig. 2g-j**) corroborated the in vivo MRI results (**Fig. 2d, e**), showing that CM injection resulted in predominant tracer accumulation in the cervical segment, whereas intrathecal injection led to an inverse rostro-caudal distribution. Additionally, ex vivo analysis demonstrated that CM injection favoured tracer dispersion in the white matter at cervical and thoracic levels, whereas intrathecal injection evoked tracer dispersion into the grey matter of thoracic and lumbar levels.

### Glymphatic Influx Occurs Preferentially along Arteries in the Spinal Cord

Previous studies have demonstrated that glymphatic flow in the brain is polarized, following a unidirectional artery-to-vein pathway with CSF influx along periarterial spaces and interstitial fluid efflux primarily along perivenous spaces and cranial nerves (Iliff et al., 2012, 2013; Jessen et al., 2015; Holstein-Rønsbo et al., 2023; Kaag Rasmussen et al., 2024; Falkenberg-Jensen et al., 2026), with a similar directional pattern reported in the optic nerve (Wang et al., 2020; Delle et al., 2024). However, the organization of tracer influx and this directional transport pattern have not yet been characterized in the spinal cord. However, this directional transport pattern has not yet been characterized in the spinal cord. To investigate this, we used NG2-DsRed mice (8–10 weeks old), in which arterioles are labelled due to DsRed expression in vascular smooth muscle, with weak to no DsRed signal in the veins and venules due to sparse or absent of smooth muscles (Illif et al, 2012; Wang et al, 2020). To focus on the early phase of CSF influx, a short circulation time of 20 min was used.

BSA^647^ was infused into CM of NG2 DsRed mice under KX anesthesia (**Fig. 3a**). 15 min after infusion, lectin (wheat germ agglutinin-488, WGA^488^) was administered intravenously via the femoral vein to label the vasculature. At 20 min post-infusion, mice were perfused with 4% PFA, spinal cords were sectioned and imaged (**Fig. 3a, b and Supplementary Fig. 3**). Confocal imaging (**Fig. 3b)** revealed robust WGA^488^ labelling (green) of nearly all blood vessels, DsRed signal in arteries and arterioles (red), and BSA^647^ tracer (fire) localized predominantly to perivascular spaces surrounding arteries (indicated by white arrows). In contrast, veins (yellow arrows) showed little to no BSA^647^ labelling, indicating minimal tracer influx along venous pathways. Segmental quantification of vessels using violin plots showed the highest vessel density in the cervical, with fewer vessels in the lumbar and the lowest density in the thoracic segment (**Fig. 3c**). Across all regions, the majority of vessels with BSA^647^ signal were arteries, followed by veins and a small subset of unclassified vessels. 65.71% of labeled vessels in the cervical regions were aeteries, and 30.37% were veins with 3.92% unclassified (**Fig. 3d and Supplementary Fig. 3**). These data indicate that spinal glymphatic influx predominantly occurs along arterial pathways, consistent with the previously described glymphatic architecture in the brain (Iliff et al, 2013), albeit less strict than the brain of which ∼90% of tracer influx occurred along arteries at (Holstein-Ronsbo et al. 2023, Fig. 3).

**Fig. 3.**
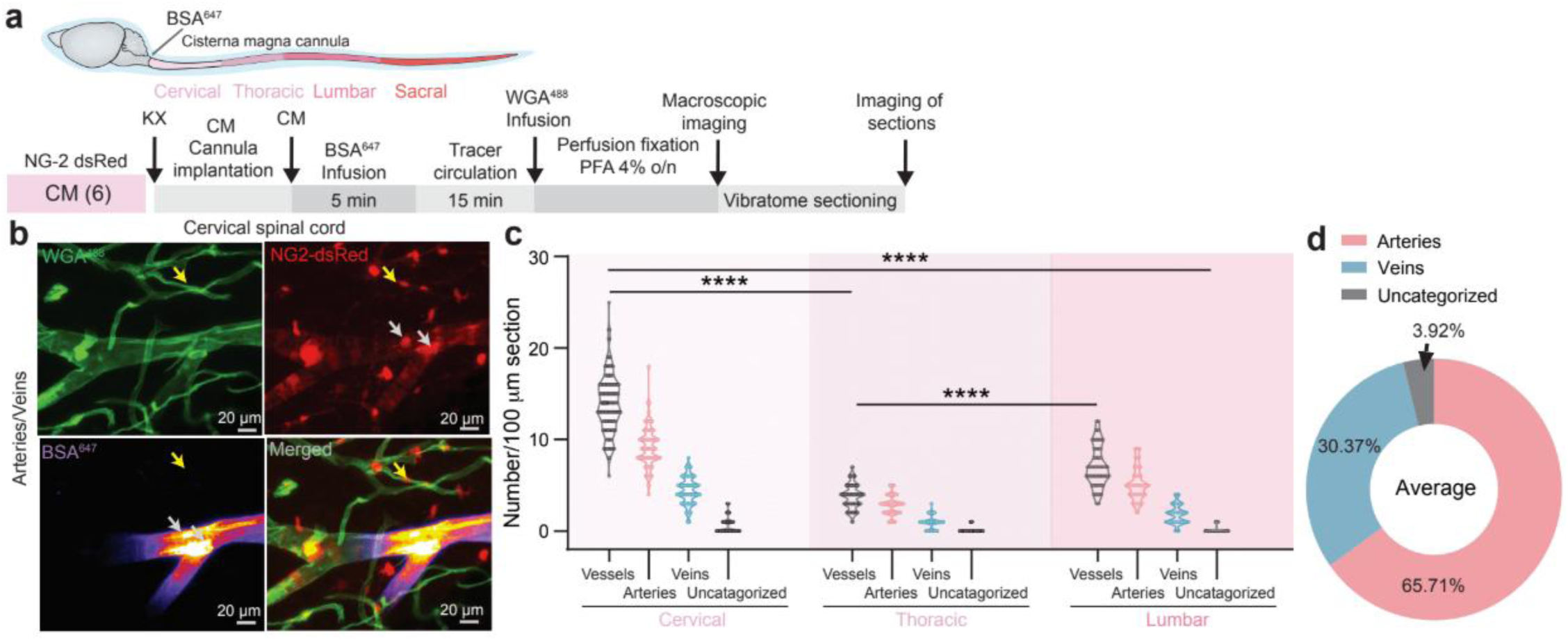
Glymphatic influx predominantly occurs along spinal arteries. Schematic of experimental methodology where bovine serum albumin conjugated with Alexa Fluor 647 (BSA^647^, 10 µl, 2 µl/min, 66.5 KDa) injected into NG2-dsRed mice via CM, followed by intravenous injection of lectin (wheat germ agglutinin, WGA^488^ conjugated with Alexa Fluor 488, green, 0.1 ml, 20 µl/sec, 38 KDa). Spinal cords were fixed, sectioned and imaged. **(b)** Confocal images (40X magnification; scale bar = 20 µm) of arteries and veins of cervical spinal cord. (**c)** Violin plots compare vessels distribution across spinal regions (Welch t-test, n=6, 14-20 sections/mouse; cervical vs thoracic: t245.9 = 36.3; cervical vs lumbar: t174.5 = 19.28; thoracic vs lumbar: t107.5 = 10.74; P ≤ 0.0001; One Way Welch ANOVA test: n=6, P ≤ 0.0001 for CM injection). **(d)** A donut plot summarizing all spinal regions (cervical, thoracic, and lumbar). Note that n is the number of mice used.

### Glymphatic Influx is Dependent on the Tracer Size

Given that MRI and ex vivo imaging data demonstrated greater contrast agent and tracer dispersion within the spinal cord following intrathecal injection, subsequent experiments were conducted exclusively via intrathecal tracer delivery. Previous studies have demonstrated that the CSF tracer influx into the brain declines with increased molecular weight; in other words, larger molecules penetrate the parenchyma to a lesser extent than smaller ones (Ichimura et al. 1991; Proescholdt et al, 2000; Abbott, 2004; Iliff et al; 2012). To evaluate whether the spinal cord also exhibits a similar size-dependent influx pattern, we compared the distribution of fluorescent tracers with different molecular weights. Specifically, OVA^647^ (45 KDa) and fibrinogen^647^ (340 KDa) were delivered intrathecally, and spinal cords were harvested 20 min later for ex vivo imaging (**Fig. 4a, b**).

**Fig. 4.**
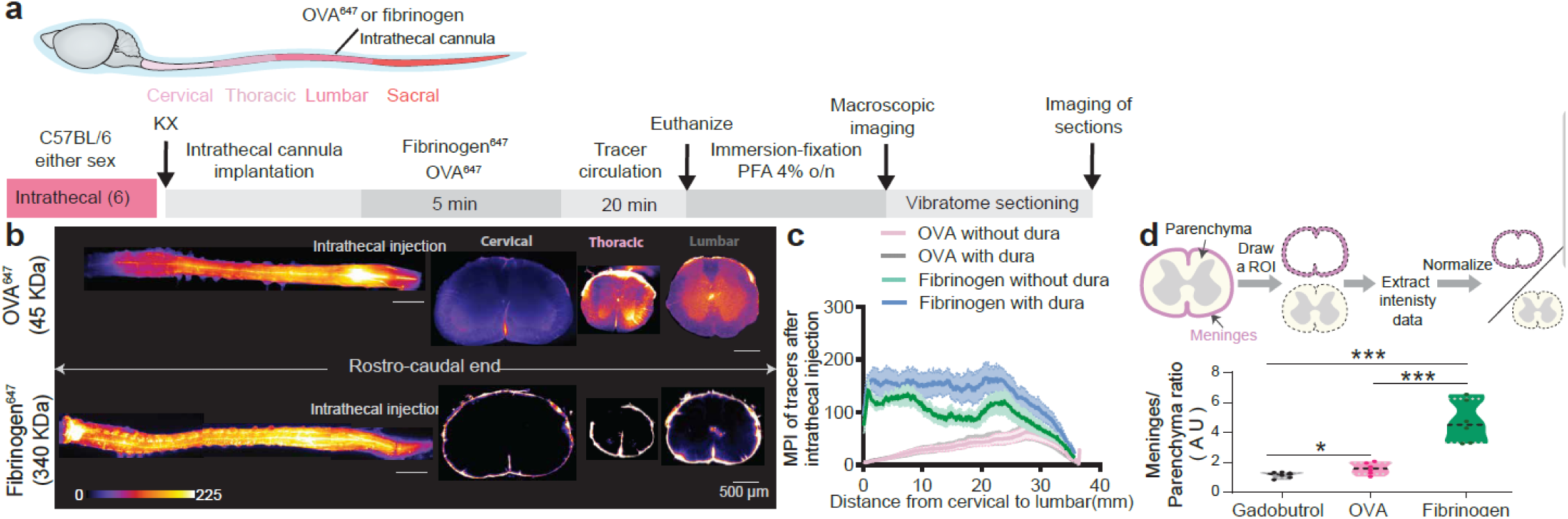
Tracer influx into the spinal cord is dependent on molecular size. **a)** Schematic of ovalbumin, OVA^647^(45 KDa), or fibrinogen^647^ (tracers conjugated with Alexa Fluor 647, 10 µl, 2 µl/min, 340 KDa) injection and extraction of spinal cords 20 min later. (**b, c)** Representative images and Mean pixel intensity (MPI) plots of OVA^647^ and fibrinogen^647^ dispersion along the spinal cord after intrathecal injection (n=6; cervical: One Way ANOVA test; P = 0.0001; thoracic: P = 0.0001; lumbar: P = 0.0001). **d)** A sketch showing meninges and parenchymal area of the spinal cord and normalization of the intensity values plotted as a violin plot (gadobutrol vs OVA^647^, P=0.05; OVA^647^ vs fibrinogen^647^, gadobutrol vs fibrinogen^647^, P<0.0001; t-test; n=6). Note that n is the number of mice used.

Representative images in **Fig. 4b** exemplify limited distribution of OVA^647^ in both the meninges and parenchyma of the cervical spinal cord. In contrast, fibrinogen^647^ was restricted to the meninges, with no detectable penetration into the parenchyma. In the thoracic region, OVA^647^ showed moderate dispersion into both meninges and parenchyma, whereas fibrinogen^647^ remained confined to the meningeal space. In the lumbar region, OVA^647^ exhibited maximal distribution in both meninges and parenchyma, while fibrinogen^647^ was still restricted to the meninges (**Fig. 4b**). Line plots in **Fig. 4c** further illustrate this pattern: OVA^647^ distribution was lowest in the cervical and highest in the lumbar region. Fibrinogen^647^ intensity, however, remained localized to the meninges across all spinal segments, regardless of the presence or absence of the dura. Although fibrinogen^647^ signal was strong in the meninges (**Fig. 4d**), it exhibited minimal parenchymal penetration, indicating that its large molecular size prevented it from crossing the meningeal barrier.

To compare fluorescence imaging with MRI, we assessed the distribution of the fluorescent tracer alongside gadobutrol, a low-molecular-weight MRI contrast agent (605 Da), as shown in Figs. 1, 2, and 4d. Tracer distributions were normalized to enable cross-modality comparisons and are presented as violin plots (**Fig. 4d**). Following intrathecal delivery, gadobutrol was limited in the cervical, minimal in the thoracic and maximum in the lumbar segment, where it was also present in the meninges.

Together, these results confirm that tracer size is a critical determinant of spinal glymphatic influx, consistent with prior brain studies (Yang et al, 2013; Iliff and Simon, 2019; Ayyappan et al, 2026). Small-to-medium-sized tracers (OVA^647^ and gadobutrol) readily dispersed into the parenchyma, whereas larger molecules (fibrinogen^647^) were excluded and confined to the meningeal spaces (**Fig. 4a–d**).

### Clearance of an Intraparenchymally Injected Tracer from the Spinal Cord

To map the temporal characteristics of spinal tracer efflux, we bypassed CSF influx by injecting OVA^647^ (1% w/v in aCSF, 0.3 µl/, 0.1 µl/min). Segment T10-T11 injection was chosen for intraparenchymal injection and the T4 segment was selected to show spinal nerves intraparenchymally. The spinal cords were extracted at 1, 8, and 24 h post-injection, followed by immersion-fixation and imaging (**Fig. 5a, b**). Tracer was confined to the ipsilateral spinal cord at 1 h, reached maximal spread with midline crossing by 8 h, and was largely cleared by 24 h. A spinal sketch in **Fig. 5c** displays the injection site and data analysis strategy. Line plot analysis (**Fig. 5c**) showed initial tracer confinement to the injection site, with gradual dispersion to the contralateral side by 8 h, and near-complete clearance by 24 h. Tracer kinetics differed between dorsal and ventral compartments: tracer levels remained stable from 1 to 8 h before a sharp decline at 24 h, whereas, ventral regions showed a significant increase at 8 h followed by a similar decline at 24 h (**Fig. 5c**). MPI remained higher at the injection than the contralateral side at both 1 and 8 h, before equalizing at 24 h (**Fig. 5d**) with a similar pattern observed dorso-ventrally (**Fig. 5e**). Tracer dispersion area also followed this temporal profile: minimal at 1 h, maximal at 8 h, and reduced by 24 h (**Fig. 5f, g**) with a greater spread on the injection/dorsal side. Transverse and longitudinal spread increased from 1 to 8 h and decreased significantly by 24 h (**Fig. 5h**).

**Fig. 5.**
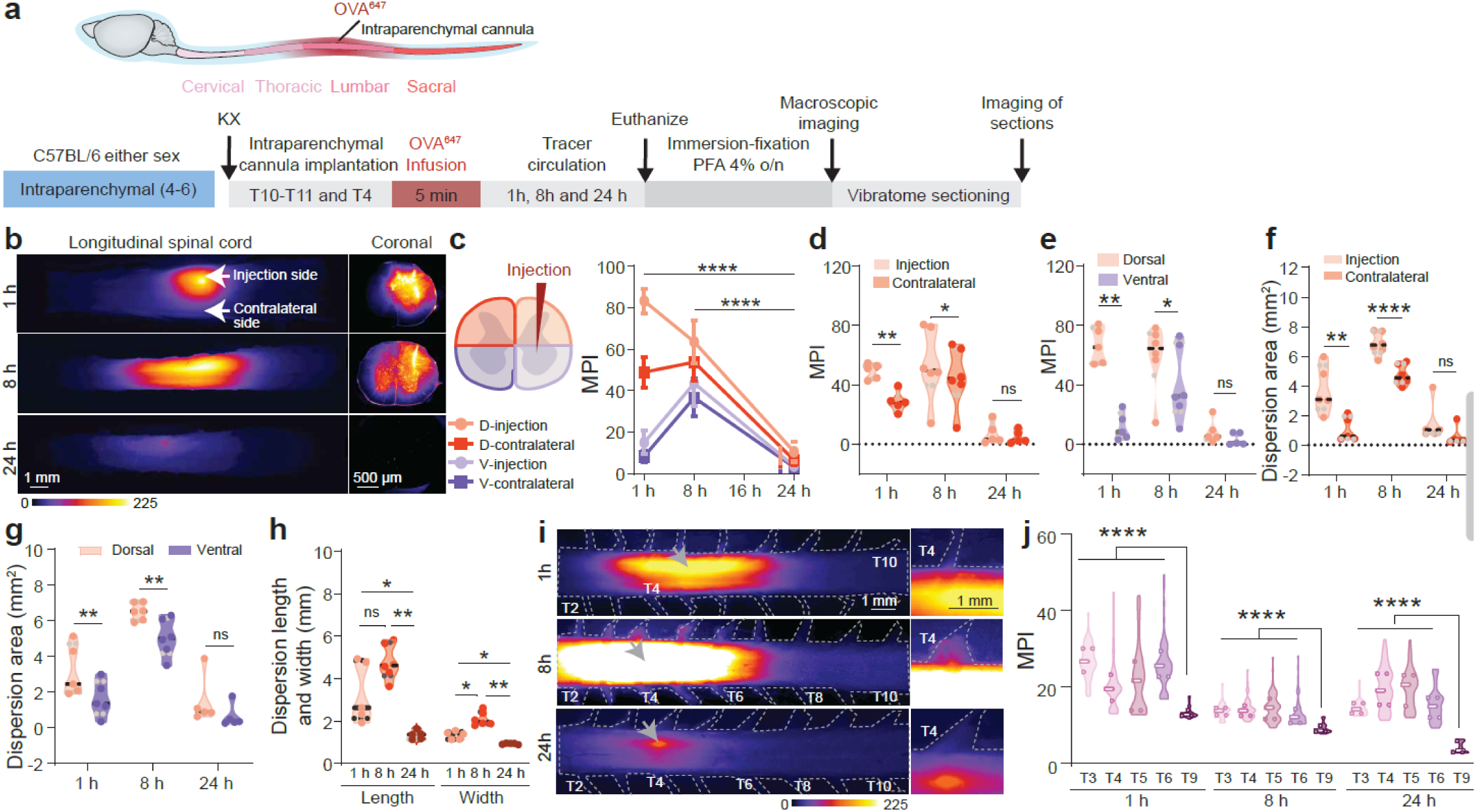
Fast glymphatic tracer clearance in the spinal cord following intraparenchymal injection. **(a)** Schematic of the experimental timeline: 5 min intraparenchymal injection of ovalbumin conjugated with Alexa Fluor 647 (OVA^647^) followed by tissue collection at 1, 8, or 24 h. **(b)** Representative images of thoraco-lumbar spinal cord whole mounts and sections at each time point showing tracer distribution. **(c)** Spinal cord sketch illustrating the strategy where D represent dorsal, and V represents ventral spinal cord. Line plot analysis quantifying pixel intensity across dorso-ventral axis (t-test, 8h vs 24 h, P=0.0001, n=4-6). **(d, e)** Quantification of MPI at 1 h, 8 h and 24 h (injection vs contralateral side: 1h, P=0.01, 8h, P=0.05; dorsal vs ventral side: 1h, P=0.01, 8h, P=0.05; t-test; n=4-6). **(f, g)** Dispersion area (8h: injection vs contralateral side, P=0.0001; dorsal vs ventral side, P=0.01; t-test; n=4-6). **(h)** Quantification of tracer spread in transverse and longitudinal direction (length: 1 h vs 24 h: P=0.05; 8 h vs 24 h, P=0.01; Width: 1 vs 24h, P=0.05; 8 vs 24 h, P=0.01; 1 vs 8 h, P=0.05; t-test; n=4-6). **(i, j)** Macroscopic images of thoracic spinal nerves (T2-T10) with injection site shown by arrows and the MPI was measured in the spinal nerves at different time points (two-way ANOVA; 1h, 8h, 24h; P=0.0001, n=4-6, N=8-10) NOTE: n is the number of mice used, and N is the number of spinal cord sections analyzed per mice.

Fluorescent imaging of spinal nerves, where the fluorescent signal from only the spinal nerves was measured, showed a strong signal between T3-T6 near the T4 injection site compared to distant levels (T9), particularly at early time points (**Fig. 5i, j**). Total tracer intensity in thoracic nerves was highest at 1 h and lowest at 24 h (**Fig. 5i, j**), suggesting slow but extended efflux at 24 h.

In summary, these data reveal that a medium intraparenchymal spinal tracer (45 KDa) is mostly cleared by 24 h post-injection and at least in part leaves by efflux along spinal nerves.

### Impaired Glymphatic Influx Dynamics after SCI

To investigate spinal glymphatic dynamics following SCI, while minimizing disruption to the dura mater and vertebral cavity, we employed a minimally invasive SCI model based on localized intraparenchymal administration of N-methyl-D-aspartate (NMDA, 100 mM, 0.3 µl) (Leem et al., 2010) at the T11 vertebra (L1 spinal level) (**Fig. 6a**) . Importantly, this approach preserved the dura and surrounding bony structures, thereby maintaining the integrity of CSF transport pathways.

**Fig. 6.**
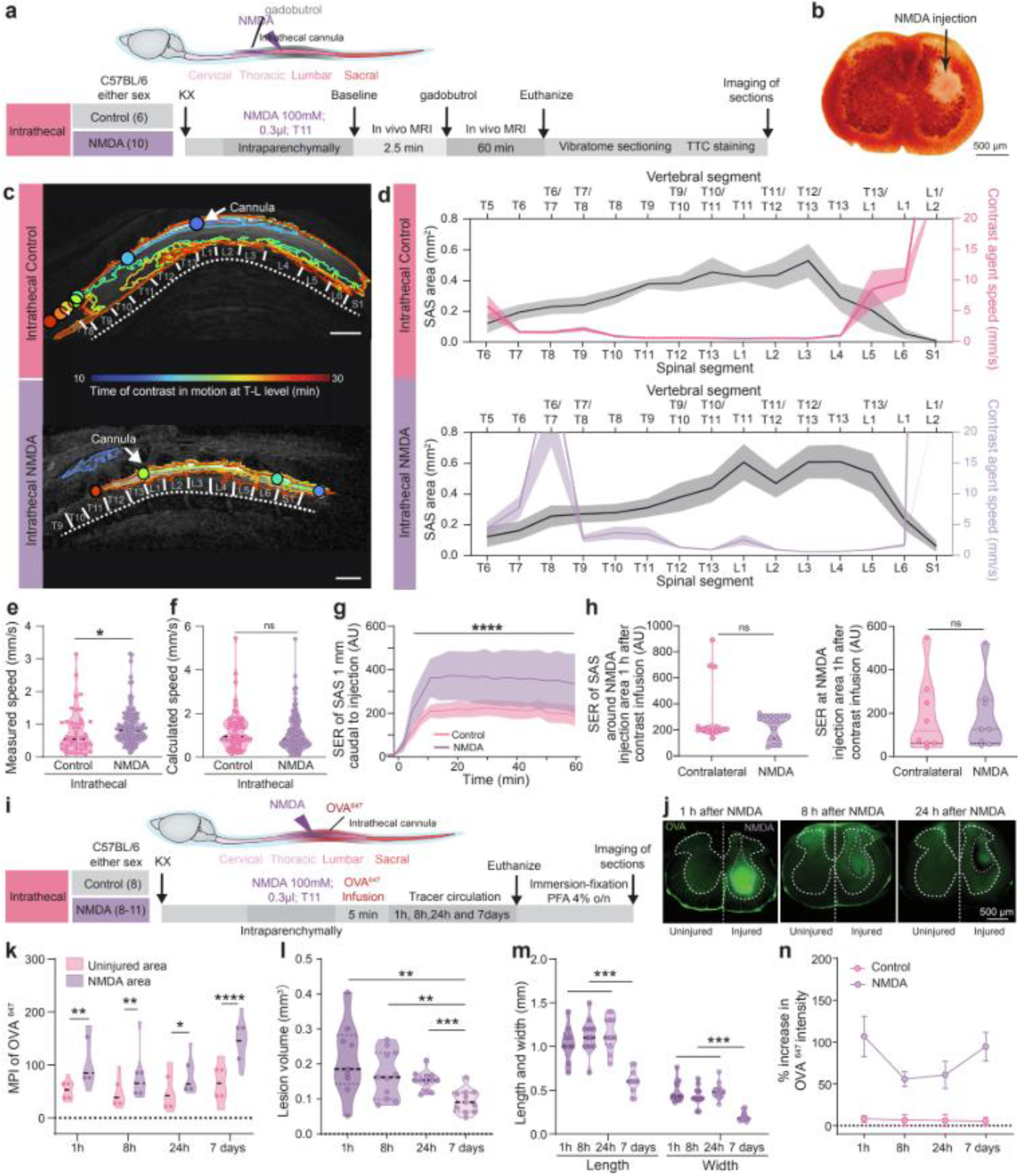
Impaired glymphatic dynamics after NMDA-induced SCI. a) Experimental design included injection of NMDA (100 mM, 0.3 µL, 0.1 µl/min, dissolved in 0.1 M PBS) at the T11 level and 24 h after 2.5 min baseline acquisition and dynamic contrast-enhanced (DCE) MRI for the next 60 min. Mice were euthanized and TTC staining followed by sectioning and imaging. **b)** TTC staining. **c)** Representative images of intrathecal control and NMDA groups illustrating the methodology to calculate contrast speed. Circles mark locations where speeds came from direct measurements. Scale bar = 2mm **d)** Contrast agent speed in the spinal SAS together with SAS area for intrathecal control and NMDA (control: n=5; NMDA: n=9; 2way ANOVA; speed: P<0,0001 when comparing various spinal levels, P=0.0029 for control vs NMDA; area: P<0,0001 when comparing various spinal levels, P=0.0334 for control vs NMDA). **e, f)** Measured (two-sample t-test, n = 6 intrathecal control, n = 10 intrathecal SCI, P = 0.035) and calculated flow speeds (tracer displacement rates) of the contrast agent in the spinal subarachnoid space (SAS) during control and NMDA intrathecal injection. **g)** Signal enhancement ratio (SER) of the contrast agent in SAS (n=5, Mann-Whitney test, P<0,0001). **h)** SER of SAS 1 h after the contrast injection around NMDA versus contralateral side 24h after NMDA injection (n = 5; 2-way ANOVA; Control vs NMDA, P<0.0001). SER (1 h after contrast infusion) of NMDA injection site versus contralateral site 24 h after lesion (n=5). **i)** Schematic illustrates intraparenchymal injection of PBS or NMDA at T11 segment in 4 groups of mice. Intrathecal tracer, ovalbumin conjugated with Alexa fluor 647 (OVA^647^, 10 µl, 2 µl/min) was administered at various post-injury time points and spinal cords were collected at 1 h, 8 h, 24 h, and 7 days after injury, fixed, sectioned, and imaged. After that MPI of OVA^647^, lesion volume, it length and width and % increase in intensity of OVA^647^ were calculated. **j)** Microscopic imaging of transverse lumbar spinal sections at 1 h, 8 h, and 24 h after NMDA injection showing NMDA injection site and contralateral site. Scale bar = 500 µm. **k)** Mean pixel intensity (MPI) of OVA^647^ at the injured site versus contralateral side in NMDA group (1 h: P = 0.0023, 8 h: P = 0.0056, 24 h: P = 0.0093, 7 days: P < 0.0001; paired t-test; n = 8–10). **l)** Tracer-labeled lesion volume (1 h vs. 7 days: P = 0.0022; 8 h vs. 7 days: P = 0.0035; 24 h vs. 7 days: P = 0.0002; t-test; n = 8–10). **m)** Tracer diffusion distances (axial and rostro-caudal) (One-way ANOVA, n = 8–10; 7 d vs. 1, 8, 24 h: all P < 0.0001). **n)** Percentage tracer intensity in control versus NMDA groups (two-way ANOVA, P=0.0001, n=9-11).

At 24 h post-injury, in vivo DCE-MRI was conducted in anesthetized mice, with baseline acquisition for 2.5 min, followed by intrathecal contrast delivery and 60 min monitoring (**Fig. 6a, c–h**). NMDA-induced SCI produced a localized sharply defined necrosis and transient paralysis. Representative staining with 2,3,5-triphenyltetrazolium chloride (TTC) 24 h post-injection confirmed a local, extensive and sharply defined necrotic lesion (**Fig. 6b**), consistent with previous reports (Faulkner et al., 2004; Leem et al., 2010).

MRI analysis revealed that after injury and later during intrathecal delivery, the contrast arrived the rostral SAS at T13 spinal level within 6.0 min and the caudal SAS at L2 level within 4.8±0.66 min; the white matter rostrally within 6.0±1.47 min at T13 and caudally within 6.0±1.2 min at L2 level; and the rostral grey matter within 6.0±1.69 min at T13 and caudal grey matter within 5.4±1.0 min at L2 level (**Supplementary Fig. 4a-d).** The temporal profile of intrathecal contrast arrival remained similar before and after NMDA injection (**Supplementary Fig. 4b-d**).

To explore if the contrast influx from SAS into the grey or white matter is dependent on the tissue anatomy and later determine whether influx changes after injury, we measured the lag time. The lag time from SAS entry to white and grey matter penetration was similar which was approximately 0.9±0.17 min and 0.93±0.16 min, respectively (**Supplementary Fig. 4e**). Thus, there was no significant difference in the white matter lag time, whereas grey matter lag time was substantially shorter after the SCI in comparison to control (**Supplementary Fig. 1e, f** and 4e). A salient observation from the intrathecal experiments was the accumulation of greater amounts of contrast agent in NMDA mice relative to controls, as demonstrated by arrival time heatmaps (**Supplementary Fig. 1a vs 4a**). As intrathecal contrast delivery showed greater spread of contrast along the thoraco-lumbar regions (**Fig. 1b, c and 2**) and the NMDA-induced lesion site was at present at the T11 level of the spinal cord (**Fig. 6**), subsequent experiments were conducted using the intrathecal delivery route. The time of contrast agent in motion was computed (**Fig. 6c)** by using the same methodology as shown in **Fig 1e**. Furthermore, the SAS area and contrast speed were compared between control and NMDA-injected mice, with contrast delivered intrathecally in both groups. Compared with intrathecal contrast delivery in control mice, NMDA-injected mice (**Fig. 6d**) showed a higher peak speed at the T7–T8 levels (Control: 1.53 ± 0.23 mm/s; NMDA: 19.18 ± 7.5 mm/s). However, the speed began to decrease at T9 in NMDA-injected mice and was minimal at L2–L5. In comparison, contrast propagation was drastically increased at the S1 segment in NMDA-injected mice relative to controls (S1 segment; Control: 42 ± 17.9 mm/s; NMDA: 182 ± 70.22 mm/s) (**Supplementary Fig. 5, showing individual mouse data**). The overall mean SAS area (Control: 0.26 ± 0.05 mm²; NMDA: 0.36 ± 0.08 mm²) and contrast speed (Control: 5.17 ± 1.7 mm/s; NMDA: 19.2 ± 7.3 mm/s) were both higher in NMDA-lesioned mice than in controls. This indicates that a focal NMDA-induced lesion is sufficient to produce measurable, segment-specific changes in both SAS geometry and contrast propagation speed at sites distant from the injury epicenter, highlighting the sensitivity of DCE-MRI-based contrast tracking as a tool for detecting altered SAS dynamics after SCI. When the measured and calculated speed (see methods section for details, **Fig. 6e, f**) were compared between intrathecal control and NMDA conditions, contrast propagation speed, specifically the measured speed (**Fig. 6e**), was accelerated following spinal lesion, consistent with the significantly greater SAS SER observed at a site 1 mm caudal to the injection over the 60 min (**Fig. 6g**). However, when SER was measured within the SAS at the NMDA injection site, spanning the T13, L1, and L2 segments, 24 h after injection, no enhanced gadobutrol influx was observed on either the NMDA-injected or contralateral side, even 1 h after contrast infusion (**Fig. 6h, left graph**). Similarly, the SER measured within the grey matter at the NMDA injection site showed no enhancement with respect to the contralateral site at 24 h post-lesion (**Fig. 6h, right graph**). Collectively, these analyses indicate that contrast propagation speed, specifically the measured speed, within the SAS was accelerated following spinal lesion. However, no enhancement in contrast influx 24 h after the lesion points to preservation of tissue resistance.

To further examine glymphatic dynamics post-NMDA injection, OVA^647^ was delivered intrathecally, and spinal cords were harvested at 1 h, 8 h, 24 h, and 7 days post-lesion (**Fig. 6i**). Fluorescence intensity was quantified in fixed spinal sections at the lesion and contralateral side (**Fig. 6j**) and tracer accumulation at the injured site at 1 h, 8 h, and 24 h. Consistently, violin plots demonstrated elevated MPI at the lesion relative to the contralateral side from 1 h to 7 days post-lesion (**Fig. 6k**).

Tracer dispersion volume at the lesion site was significantly reduced at 7 days post-lesion relative to earlier time points (1h, 8h and 24 h), with no difference in 1-24 h (**Fig. 6l**). To assess the extent of tracer penetration into parenchyma, axial and rostro-caudal diffusion distances from the lesion site were quantified (**Fig. 6m**). The tracer dispersed across 1-2 spinal levels from the lesion (1.352±0.127 mm) with a significant reduction in dispersion distance in both directions 7 days post-injury, although no significant differences were detected among the 1 h, 8 h, and 24 h groups (**Fig. 6m**). Relative tracer intensity was further calculated as a % increase over background fluorescence at the lesion site (**Fig. 6n**). Mean % increases were 106.7±24.2 at 1 h, 55.6±9.1 at 8 h, 60.7±16.3 at 24 h, and 94.5±17.1 at 7 days post-SCI, with no significant differences across time points within the same condition (**Fig. 6n**).

Collectively, these results demonstrate that intrathecal contrast administration following SCI accelerated glymphatic flow and enhanced tracer accumulation within the spinal parenchyma, suggesting impaired clearance and potential glymphatic dysfunction.

### AQP4 Deletion Suppressed Spinal Glymphatic Dynamics and Aggravated Injury

Aquaporin-4 (AQP4) is a water channel expressed in the astrocytic endfeet that facilitates CSF inflow along perivascular spaces, and its genetic deletion or pharmacological inhibition reduces glymphatic flow by 25–70% (Iliff et al, 2012, Mestre et al, 2018; Gomolka et al, 2023; Giannetto et al 2024). SCI studies have reported spinal edema increases after the injury in parallel with increased AQP4 expression (Nesic et al, 2006; Li et al, 2018), while AQP4 deletion reduces edema (Saadoun et al, 2008; Huang et al, 2019; Garcia et al., 2023). Pan et al. (2019) In contrast, other reports show that AQP4 deletion aggravated edema, caused neuronal loss, and impaired motor recovery in a mouse model of contusion SCI (Kimura et al, 2010). Here, we compared wild-type C57BL6 (C57-PBS), C57BL6 NMDA-injured (C57-NMDA) and AQP4 knockout C57BL6 injured (AQP4KO-NMDA) mice.

We first compared hindlimbs motor functions, daily for 1 week before and after SCI (**Fig. 7a**), using the Basso Mouse Scale (BMS, Basso et al., 2006). BMS score was normal (marked 9) before the lesion and dropped to 0 at day 0 in injured groups as no ankle movement was observed (**Fig. 7b)**. Afterwards, the BMS score improved daily but to a variable degree in the different groups. In C57-PBS mice, all the mice recovered to a score 9 after day 0. C57-NMDA mice showed a modest score increase each day until day 7 after the injury, with a significant motor recovery on day 7 compared to day 1-2. However, AQP4KO-NMDA mice showed no or limited improvement in BMS score on day 1-2. Interestingly by day 7, BMS score of all groups was significantly improved compared to day 0.

**Fig. 7.**
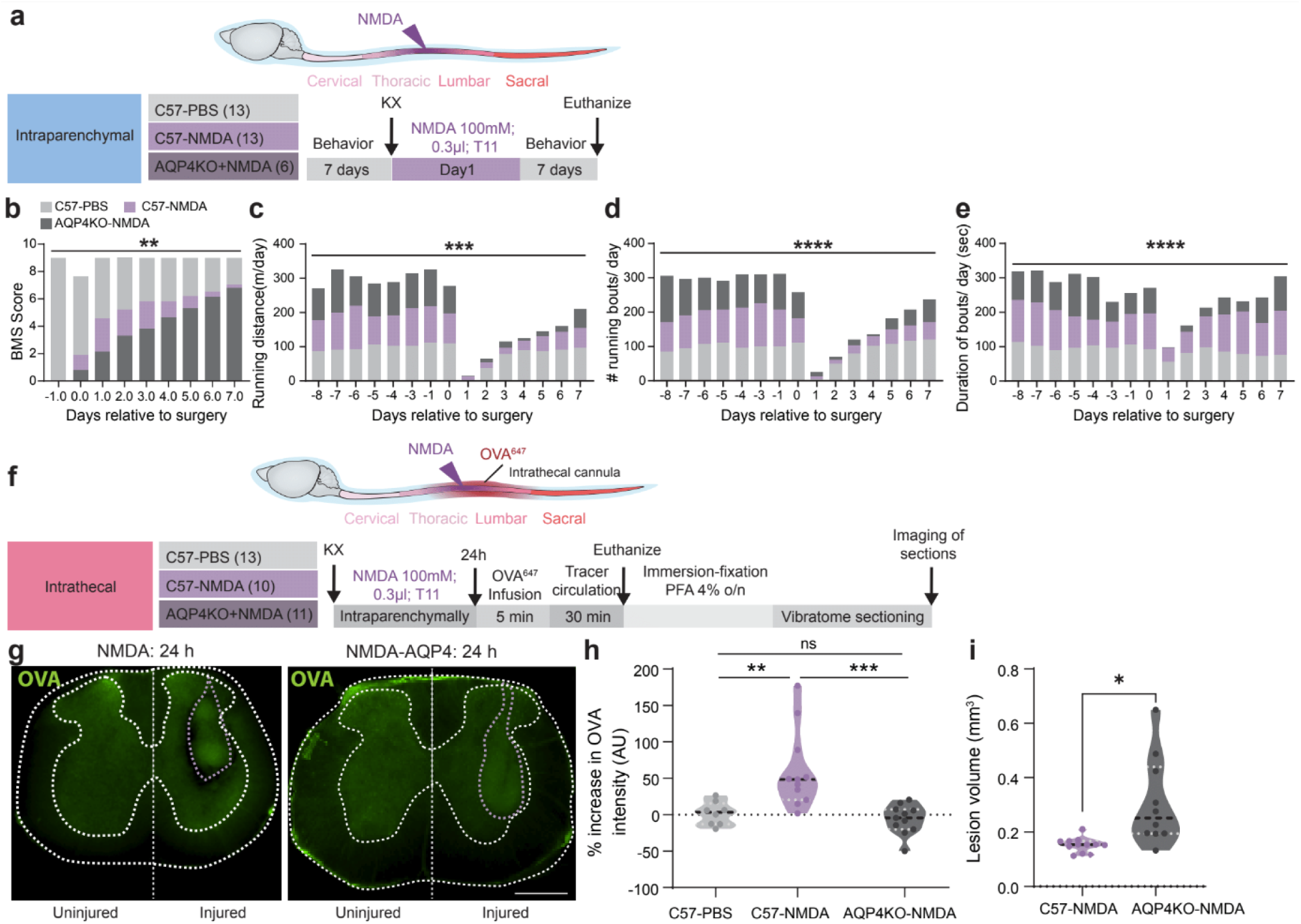
Impaired glymphatic dynamics after NMDA-induced SCI. a) Behavior paradigm including Basso mouse scale (BMS) score and voluntary wheel test before and after the NMDA injection administered intraparenchymally (100 mM, 0.3 µL, 0.1 µl/min, dissolved in 0.1 M PBS) in three groups of mice: C57BL6 wildtype mice with PBS injection (C57-PBS), C57BL6 wildtype mice with NMDA injection (C57-NMDA) and C57BL6 AQP4KO mice with NMDA injection (AQP4KO-NMDA). **b)** BMS scores before and after NMDA-induced SCI (Two-Way ANOVA test: P=0.0096; t-test: day 0 vs day 7; C57-NMDA (n=13): 0.0001; and AQP4KO-NMDA(n=6): 0.0002). n for C57-PBS mice = 13. **c-e)** In voluntary wheel exercise test, stacked histograms depicts running distance (day 1 and day 7: Two-Way ANOVA test: P=0.0005, t-test: P = 0.0134), number of the running bouts/day (Two-Way ANOVA test: P=0.0001, t-test: P = 0.0001) and duration of running bouts/day (AQP4KO-NMDAvs C57-PBS: t-test, P=0.0195) after NMDA induced-SCI. **f)** Scheme demonstrating intrathecal injection of ovalbumin conjugated with Alexa fluor 647 (OVA^647^) and its circulation for 24 h in three groups of mice: C57-PBS, C57-NMDA and AQP4KO-NMDA. **g)** Images showing lesioned spinal cord samples from C57-NMDA and AQP4KO-NMDA mice 24 h after the injury. Scale bar = 500 µm **h)** Percentage increase in intensity of OVA^647^ (C57-PBS vs C57-NMDA: P=0.01; C57-NMDA vs AQP4KO-NMDA: P=0.001; C57-PBS, n=13, C57-NMDA, n=10, AQP4KO-NMDA, n=11). **i)** Lesion volume based on OVA^647^ distribition patterns(t-test, P = 0.0117).

Further, we used a voluntary wheel exercise test to promote and assess the recovery of motor function and mice were trained prior to injury. Here, we measured the total running distance, the number and duration of individual bouts each day before and after the SCI. We observed almost a complete loss of ability to perform the running wheel test on 1 day and 2 after the lesion followed by slow recovery during the 7 days (**Fig. 7c-e**). During 7 days after the lesion, the mean running distance, the mean number of running bouts and the mean duration of running bout per day were notably lower in the C57-NMDA and AQP4KO-NMDA mice than the control (C57-PBS) mice. However, 7 days after the lesion, C57-NMDA mice running distance was improved between day 1 and 7, with no significant difference in the running distance and number of bouts in AQP4KO-NMDA mice (**Fig. 7c-d**). When the duration of running bouts/day was measured, AQP4KO-NMDA mice ran for a shorter duration at day 1 after the lesion; however, other groups ran for a longer duration, with recovered bouts duration in all groups at day 7 (**Fig. 7e**). The data may indicate that C57-NMDA mice improved motor recovery, whereas AQP4KO-NMDA group did not recover significantly, suggesting persistent motor impairment.

Further to unveil the extent of spinal lesion and changes in the glymphatic dynamics 24 h after injury, another set of mice were anesthetized, OVA^647^ was injected (2 µl/min) intrathecally, sacrificed after 30 min tracer circulation, followed by immersion-fixation and vibratome sectioning (**Fig. 7f**). **Fig. 7g** depicts that 30 min after injecting OVA^647^ into injured mice, C57-NMDA mice showed smaller lesion area whereas AQP4KO-NMDA mice exhibited a larger lesion area. In lumbar spinal cords, % intensity of OVA^647^ was higher in NMDA mice than both C57-PBS and AQP4KO-NMDA mice (**Fig. 7h**). When their lesion volume was computed by Luxol blue staining, AQP4KO-NMDA mice exhibited increased lesion size compared to C57-NMDA mice (**Fig. 7i**).

These results suggest that, like in the brain, AQP4 plays a key role in spinal glymphatic influx. Its deletion suppresses glymphatic function and exacerbates spinal lesion, as evidenced by worsened hindlimb motor function and increased lesion volume (Wu et al, 2014; Garcia et al, 2023).

## Discussion

The present study provides the first comprehensive in vivo characterisation of the CSF and glymphatic dynamics along the mouse spinal cord and demonstrates the major bottlenecks in the SAS present in the mid-to-lower cervical (C3-C7), mid-to-lower thoracic (T6-T12) and lower lumbosacral (L5-S1) levels, where the speed of the contrast agent increased abruptly. Further, using in vivo MRI and ex vivo fluorescent imaging strategies, this study exemplifies the influx of contrast and fluorescent tracer in the white and grey matter when injected via CM or intrathecally. Surprisingly, the dispersion of the contrast agent was most extensive in the cervical and thoracic regions when injected via CM, whereas its dispersion peaked in the thoraco-lumbar regions when injected intrathecally. Moreover, more tracer accumulated in white matter when the contrast agent was injected via CM. In contrast, intrathecal injection led to higher tracer dispersion in grey matter regions. We further show that glymphatic influx in the spinal cord, similar to the brain, occurred primarily along arteries and was dependent on the molecular size of the tracer. Moreover, tracers injected into the spinal tissue efflux, at least in part, along spinal nerves. When a focal spinal lesion was induced by a local NMDA injection, increased CSF-measured speed was observed in the SAS, especially at mid-thoracic (T7-T8) and upper sacral (S1) levels, and increased retention of tracer at the injured area was noted. Furthermore, genetic deletion of AQP4 not only impaired motor recovery after SCI but also suppressed glymphatic function and increased lesion volume. Together, these findings indicate a functional glymphatic system in the spinal cord and that AQP4-dependent fluid transport plays a role in the secondary injury progression.

A 3D computational model study showed that CSF velocities varied across anatomical regions of the human SAS (Linge et al, 2010). Similarly, phase-contrast MRI in the cynomolgus monkey spinal cord revealed an inverse relationship between SAS geometry and CSF velocity (Khani et al, 2019). However, to our knowledge, no studies have directly compared segment-wise spinal SAS area with contrast agent speed in the spinal SAS. Bessen et al. (2023) showed greater CSF flow at C2–C3 and the lowest flow at T8–T9 and L1–L2 in pigs. Despite species differences, our data showed a similar pattern, with increased contrast propagation at C3 up to C7 that decreased at T8–T10 and L1–L4. The spinal segment-wise contrast propagation speed measured in the current study is among the first segment-resolved measurements of net contrast speed reported in a rodent model, which ranges between 0.49-20.42 mm/s depending on the segment. Direct rodent comparators are scarce, reported CSF velocities in mice and rats are largely confined to the cranial compartment and are one to three orders of magnitude slower than our values, for example, 0.9–5.5 µm/s in mice brain (Bedussi et al, 2018) consistent with the general principle that CSF flow velocity scales with the size of the CSF compartment and is markedly slower in rodents than in larger animals such as pigs (0.9-1.41 m/s, Bessen et al, 2023) or humans with reported pulse wave velocity of 19.4 cm/s (Sass et al, 2017) in the spinal cord SAS. A net non-pulsatile bulk CSF motion along the spinal canal is considerably slower and is around 0.17 mm/s (Sánchez et al., 2018). Together, these comparisons highlight both limited studies on segment-wise spinal CSF transport data in mice and the importance of comparing different physical quantities such as propagation speed, pulsatile velocity and net bulk flow when comparing across studies and species. Our DCE-MRI analysis identified distinct bottleneck zones along the spinal cord, where the SAS areas along the spinal segments were minimal, and the CSF contrast speed was higher. These constrictions were located at the C3-C7, T6-T12 and L5-S1 spinal segments (**Fig. 1d**). Consistent with a Venturi-like effect (Greitz, 2006; Clarke et al., 2013), contrast movement through the SAS was fastest in the cervical (C3–C7), thoracic (T6–T12), and lower lumbar (L5–S1) regions, suggesting that local narrowing of the SAS accelerated CSF flow through these segments (**Fig. 1d**). Importantly, when contrast propagation was compared between CM and intrathecal injection routes, there was no significant difference between the two routes (**Fig. 1e, f**), indicating that this regional speed profile reflects an intrinsic property of SAS geometry rather than the site of contrast administration. This has direct implications for intrathecal drug and gene therapy delivery, where the rate and extent of agent spread through the SAS are key determinants of treatment efficacy. Current data suggest that regional bottlenecks, rather than the injection site, are the dominant factor shaping how quickly an intrathecally delivered agent reaches distant spinal levels. Moreover, the identification of consistent, geometry-driven flow increase at the cervical, mid-to-lower thoracic, and lower lumbosacral bottlenecks may help reveal the reason for vulnerability of these regions to secondary injury and impaired CSF clearance after trauma, as a minor narrowing at a bottleneck segment could disturb the fluid flow.

Prior work demonstrates that dispersion of contrast agent is dependent on the route of delivery (Hinderer et al, 2020; Khani et al, 2022; Jacob et al, 2022; van Osch et al, 2024). A study on primates showed that AAV delivery via lumbar puncture resulted in lower transduction into the brain and the cervical spinal cord compared to CM injection (Hinderer et al, 2020). A computational fluid dynamics study of human in silico trials showed that cisterna magna (CM) injection delivered a greater therapeutic dose to brain regions, whereas lumbar puncture/intrathecal injections could target spinal cord regions (Khani et al, 2022). Quantitative ex vivo fluorescent imaging corroborated the in vivo MRI results and again suggested that the ratio of contrast agent/tracer in white versus grey matter was strongly influenced by the route of delivery (**Fig. 2**). The contrast agent delivered via CM dispersed widely and evenly in the cervico-thoracic regions, whereas, when it was delivered intrathecally, it dispersed more in the thoraco-lumbar regions with the higher peak SER at the injection site and gradually reduced peak SER with respect to the distance (**Fig. 2 and Supplementary Fig. 1**). Furthermore, ex vivo fluorescent imaging analysis suggested that the tracer was more dispersed in the cervico-thoracic white matter when injected via CM. In contrast, it was more dispersed in the thoraco-lumbar grey matter when injected intrathecally (**Fig. 2g-j**). These differences in delivery route and the dispersion of the contrast/tracer may be particularly relevant for targeted intrathecal drug delivery strategies in SCI.

It is well established that glymphatic influx in the brain occurs predominantly along arteries (Illif et al, 2013; Holstein-Rønsbo et al, 2023), but whether a similar pathway exists in the spinal cord has not previously been investigated. Using NF2-DsRed mice, in which arterial smooth muscle cells express DsRed, we found that tracer delivered via CM entered the spinal cord primarily along arteries rather than veins at early time points (**Fig. 3 and Supplementary Fig. 3**). Approximately 65% of tracer-labeled vessels in the cervical cord were arteries, whereas venous labeling was comparatively limited. This arterial preference was observed across all spinal regions, with the highest vascular density and tracer signal detected in the cervical segment. This pattern was consistent with the arterial glymphatic influx pathway previously described in the brain (Illif et al, 2013; Mestre et al, 2018; Holstein-Rønsbo et al, 2023). Together, these findings extend the polarized artery-to-vein organization of glymphatic transport, previously demonstrated in the brain and optic nerve (Wang et al., 2020; Delle et al., 2024), to the spinal cord. The identification of a functional perivascular glymphatic influx pathway in the spinal cord has important implications for the clearance of injury-related metabolites and inflammatory mediators after SCI.

Prior studies have reported that the tracer influx is dependent on its molecular size in the brain (Ichimura et al, 1991; Yang et al, 2013; Iliff et al, 2013; Zhu et al, 2023) and the spinal cord (Lam et al, 2017). Small molecular weight contrast agent, gadobutrol (605 Da) and intermediate-sized tracer, OVA^647^ (45 KDa) penetrated the spinal parenchyma with the greatest penetration in the lumbar and thoracic regions, followed by the cervical spinal segment (**Fig. 4**). The large molecule, fibrinogen^647^ (340 KDa), on the other hand, was entirely confined to the meningeal space and showed no parenchymal penetration regardless of the spinal level. These findings imply that the spinal perivascular space imposes a size-dependent barrier to macromolecular transport, an important consideration both for understanding the normal physiological clearance capacity of the spinal glymphatic system and for designing CSF-based drug delivery strategies targeting SCI.

Solute clearance pathways following intraparenchymal brain injection of fluorescent or radiolabeled tracers have been extensively documented (Cserr et al, 1981; Iliff et al, 2012; Xie et al, 2013; Peng et al, 2016). In contrast, complete efflux pathways in the spinal cord have received limited attention, although a few studies have reported the existence of spinal efflux routes (Liu et al, 2018; 2021). Here, intraparenchymal injection of OVA^647^ into the spinal cord revealed that the large tracer (45 KDa) was cleared locally over a 24 h period. Tracer spread within the cord was maximal at 8 h and was followed by near-complete clearance by 24 h (**Fig. 5**). Clearance kinetics differed between dorsal and ventral compartments at 1 h and 8 h, with higher pixel intensity and dispersion area in the dorsal compartments indicative of reduced clearance. However, in the ventral compartment, tracer intensity was lower, and it dispersed in a smaller area at 1 h. Interestingly, tracer intensity was significantly increased at 8 h, and the tracer was dispersed in a larger area even though the dispersion in the ventral region was still smaller than the dorsal (**Fig. 5 c, e, g**). Notably, spinal nerve roots appeared to serve as major efflux routes, consistent with the established role of cranial nerves as efflux pathways in the brain (Johnston et al, 2004; Koh et al, 2005; Praetorius, 2007; Rasmussen et al., 2018; Jessen et al., 2015; Jacob et al 2022). The identification of spinal nerves as interstitial fluid efflux pathways is particularly relevant to SCI. Disruption of the dura–nerve root interface after injury may impair solute clearance and thereby contribute to secondary tissue damage through the accumulation of toxic and inflammatory mediators.

Following focal NMDA-induced SCI at T11, DCE-MRI revealed a significant acceleration of intrathecal contrast propagation at the mid-thoracic (T7-T8) and lower lumbosacral levels (L6-S1) compared with controls (**Fig. 6d, Supplementary Fig. 5**), suggesting that a minor lesion in the spinal cord can disrupt SAS fluid dynamics away from the lesion site. Berliner et al. (2019) showed increased fluid flow within spinal cord tissue after SAS obstruction by extradural constriction of the rat spinal cord. They also noted swelling and inflammation of the spinal cord due to the spinal constriction. In rats, acute tissue swelling and rostro-caudal spread of damage are already evident by 24 h post-SCI (Mihai et al, 2008), consistent with our DCE-MRI acquisition timing of 24 h after NMDA-induced lesion. Further, consistent with our data (**Fig. 6g, j, k, n**), they reported contrast enhancement at 24 h post-injury. While these studies establish that swelling and edema are already present and spreading beyond the injury epicenter within 24 h, none have directly linked this early swelling to altered contrast propagation speed in the SAS, the correlation we have demonstrated here. One explanation is that swelling narrows the segment-wise SAS levels’ proximity to and further from the injury site, leading to further constriction at the bottleneck regions (shown in Fig. 1d) thereby further accelerating contrast flow through a Venturi-like mechanism. This could mean that SCI-related changes in fluid flow are not local and reflect an interaction between the injury and the anatomical geometry of the SAS where bottleneck regions are highly sensitive to even modest swelling. This would suggest that segments coinciding with SAS constrictions may be informative sites for detecting and monitoring SCI-related fluid disturbances non-invasively, and may also represent regions at heightened risk of secondary complications such as impaired clearance after injury.

The increase in measured propagation speed – defined here as a measure of how fast the tracer front spreads through the SAS (see methods for details), in the absence of changes in calculated speed suggests that the injury may have altered the local cross-sectional area of the SAS, possibly reflecting spinal cord swelling (**Fig. 6 e, f**) which was consistent with the increased SER caudal to injection after the lesion (**Fig. 6g**). However, SER at the injection site and contralateral side remained unchanged (**Fig. 6h**), indicating that net intrathecal-to-parenchymal tracer exchange between the ipsilateral and contralateral sides was not augmented at this early time point. This is consistent with the highly localized nature of the NMDA lesion model in which surrounding tissue remains intact (Faulkner et al., 2004). Ex vivo fluorescent imaging of intrathecally delivered OVA^647^ dispersion at multiple time points after SCI showed accumulation of tracer at the lesion site, even 7 days after the lesion, relative to the contralateral site; however, the lesion volume was significantly reduced 7 days after the lesion in comparison to 1,8 and 24 h (**Fig 6i, j, l-n**) (Waight et al, 2026; Mihai et al, 2008). The persistent local accumulation of tracer occurred along with a significant reduction in lesion volume between 1–24 h and 7 days post-injury (**Fig. 6l–m**), indicating that the two processes are mechanistically distinct: whereas the necrotic lesion core undergoes progressive resolution, local fluid transport dynamics remain abnormal. We interpret this as evidence that emerging glial scarring restructures the perivascular and interstitial microenvironment in a way that impedes normal tracer clearance, even as gross tissue damage resolves (Garcia et al., 2023). Whether this persistent tracer accumulation reflects a primary glymphatic dysfunction or a structural barrier imposed by the glial scar remains an important open question for future investigation. A key finding of the present study is that genetic deletion of AQP4 suppressed glymphatic influx and worsened functional (**Fig. 7a-e**) and histological outcomes (**Fig. 7f-i**) following lesion. The role of AQP4 in SCI remains debated. Several studies have suggested that AQP4 upregulation after SCI contributes to edema formation and that AQP4 deletion reduces spinal cord water content (Saadoun et al, 2008; Huang et al, 2019; Garcia et al, 2023). In contrast, other reports indicate that loss of AQP4 can worsen vasogenic edema (Papadopoulos et al, 2004; Bloch et al, 2005). The increased lesion volume observed here in AQP4 knockout mice (**Fig. 6g, i**) may reflect exacerbated edema and impaired tissue-fluid regulation. A recent study reported a substantial reduction in glymphatic clearance of AAV vector in AQP4 knockout mice (Murlidharan et al, 2016; Cona et al, 2026). The aggravated lesion volume after injury in AQP4 knockout mice is also consistent with prior evidence that impaired glymphatic clearance permits the accumulation of excitotoxic metabolites and inflammatory mediators, thereby promoting secondary injury expansion in models of traumatic brain injury and stroke (Iliff et al., 2014; Jessen et al., 2015; Cona et al, 2026).

## Limitations of the Study

The present study has several limitations that should be considered. First, the invasive nature of the experiments required ketamine/xylazine anesthesia. Although ketamine/xylazine is widely used in studies of glymphatic dynamics, the extent to which it modulates spinal glymphatic transport remains to be fully characterized. Second, the localized NMDA-induced spinal cord lesion model preserves the dura mater and produces a focal lesion, but it does not recapitulate the complex vascular, inflammatory, and mechanical events that occur in clinical SCI. Third, the study primarily focused on outcomes up to 7 days post-lesion; therefore, the long-term consequences of altered spinal glymphatic dynamics across different SCI models remain unknown. Finally, the mechanistic relationship between spinal glymphatic dysfunction and key hallmarks of delayed secondary injury, including axonal loss, neuropathic pain, demyelination, and glial scarring, remains to be established.

## Methods

### Animals and Ethical Statement

Adult C57BL/6 male and female mice (aged 10–12 weeks) were obtained from Jackson Laboratory. Aquaporin-4 knockout (AQP4(-/-)) mice (Thane et al., 2011) were routinely crossbred with wildtype (WT) mice at the University of Copenhagen and the University of Rochester. AQP4(-/-) were generated as described by Thane et al. (2011). All mouse strains were housed under a 12-h light/dark cycle with controlled temperature (22 ± 2°C) and humidity (55±10%) and were group-housed (up to five mice per cage) with access to food and water. Animal use protocols were approved by the University of Rochester Medical Center Committee on Animal Resources (license number: 2011-021E) and the University of Copenhagen Animal Inspectorate (license numbers: 2015-15-0201-00668 and 2023-15-0201-01600). All surgical procedures were performed under aseptic conditions.

### Cisterna Magna injection

Mice were anesthetized with ketamine (100 mg/kg) and xylazine (10 mg/kg, K/X). For longitudinal measurement, supplemental ketamine (50 mg/kg) was given every 45-60 min to maintain the anesthesia whenever required. A midline neck incision was made to expose the cisterna magna (CM) (Du et al., 2022). A 30 G cannula connected to a PE-10 tube filled with artificial cerebrospinal fluid (aCSF, 126 mM NaCl, 26 mM NaHCO3, 1.25 mM NaH_2_PO_4_, 2.2 mM KCl, 2 mM CaCl_2_, 2 mM MgSO_4_, 10 mM glucose, pH 7.4) was inserted into the CM and secured with superglue and dental cement. For the MRI experiment, we used a 30G copper cannula (Nippon Tokushukan, Mfg, Tokyo, Japan) to minimize image artifacts and distortion. The PE-10 tube was attached to a Hamilton syringe connected to a Harvard Instrument Syringe Pump, and an MRI contrast agent (gadobutrol, 605 Da) was infused at 1.5 µl/min with a total volume of 12 µl whereas fluorescent tracers: ovalbumin (OVA^647^, 45 KDa), bovin serum albumin (BSA^647^, 66.5 KDa), and fibrinogen (340 KDa) was infused at 2 µl/min with a total volume of 10 µl (0.5% weight/volume in aCSF). Spinal cords were removed, fixed with 4% paraformaldehyde (PFA) and macroscopic imaging was performed on the whole and sectioned spinal cords (100 µm thick sections). The choice of the tracer was dependent on the goal of the experiment.

### Intrathecal tracer injection

Mice were anesthetized with ketamine (100 mg/kg) and xylazine (10 mg/kg). After performing a small laminectomy and exposing the lumbar dura (T13-L2 spinal segment), a 30 G cannula attached to PE-10 tubing was inserted into the subarachnoid space (SAS) at the L1 segment. The tubing was attached to a Hamilton syringe, and a fluorescent tracer either OVA^647^ or fibrinogen^647^ was injected at 2 µl/min (total volume: 10 µl) or gadobutrol was injected at 1.5 µl/min with total volume of 12 µl.

### Dynamic contrast-enhanced MRI

MRI was conducted on a 9.4 Tesla animal scanner (BioSpec 94/30 USR, Bruker BioSpin) using a cryogenically cooled quadrature-resonator transmit/receive coil (CryoProbe, Bruker). Mice were placed in an MR-compatible holder, and body temperature was maintained at 37°C. Pre- and post-contrast T1-weighted imaging (T1WI) was performed using a 3D-FLASH sequence (TR = 13 ms, TE = 2.1 ms, FA = 15°, FOV = 25.6 mm × 9.6 mm × 10 mm, Matrix = 256 × 96 × 100, Spatial resolution = 100 µm, acquisition time = 2.5 min). First 3 scans were acquired without CSF contrast agent injection (baseline). Later, a contrast agent, gadobutrol was injected either into the CM or intrathecally, and dynamic T1WI scanning was conducted for 60 min. Motion correction was performed using ANTs software, and T1WI signal enhancement ratios were calculated as percent signal change from baseline. Image analysis was done in ITK-SNAP software.

### SAS Speed Measurement of DCE-MRI Data

Because contrast propagation speed did not differ substantially between CM and intrathecal injection routes in control mice, SAS area and propagation speed data from both groups were combined into a single graph as shown in Fig 1d.

#### Measured flow speeds in subarachnoid spaces

Tracer displacement rate was used as an indicator of flow speed in the SAS. For each video showing the spread of tracer over time, we chose a brightness threshold that consistently separated regions of high tracer concentration from regions where tracer had not yet reached. We then located that brightness in two subsequent frames of the video and estimated the speed as the distance between those two locations, divided by the interval between frames. Measurements were recorded on the mid-saggital plane, in which the tracer was visible since injection, and for frame pairs where there was visible tracer movement for the given threshold. In most videos, we made several such measurements (see the Supplementary Fig. 2 and 5).

#### Calculated flow speeds in subarachnoid spaces

Each flow speed measurement (made as described above) was used to calculate flow speeds in other spinal segments. The calculation was based on the approximations that cerebrospinal fluid is incompressible and passes along the SAS without escaping. With those assumptions, conservation of fluid mass requires that the volume flow rate be uniform along the spinal SAS, so

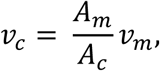

where *v_m_* is the measured speed, *A_m_* is the cross-sectional area of the spinal SAS in the segment where the speed was measured, *v_c_* is the calculated speed in another segment, and *A_c_* is the cross-sectional area of the SAS in that segment. Cross-sectional areas were determined by identifying the SAS and each spinal segment in each 3D image, calculating the volume of SAS within each spinal segment, and dividing that volume by the length of that spinal segment. Segment lengths were determined by skeletonizing the segmented spine in order to produce a centerline curve, then measuring the length of the part of the curve lying within each segment.

### Ex Vivo Fluorescence Imaging

Mice were divided into 6 major groups depending on the goal of the experiment. In group 1, OVA^647^ was injected via CM and in group 2, OVA^647^ was injected intrathecally. 120 min after the injection, mice were anesthesized with ketamine and xylazine, decapitated and spinal cords were extracted and fixed in 4% PFA. In group 3, BSA^647^ was injected via CM followed by infusion of wheat germ agglutinin (WGA, lectin) intravenously via femoral vein with 30G needle at a concentration of 1 mg/ml in 0.9% saline and intracardial perfusion was performed 20 min later with 20 ml 1X PBS followed by 4% PFA. In group 4, after the intrathecal injection of either OVA^647^ or fibrinogen^647^ tracer, spinal cords were removed 20 min later, followed by fixation in 4% PFA. In group 5, mice were anesthetized, OVA^647^ was administered intraparenchymally followed by extraction of the spinal cords at various time points such as 1h, 8h, 24h and 7 days after the tracer injection. In group 6, mice were anesthetized, and focal SCI was induced with intraparenchymal NMDA injection (10 mM, at T11). OVA^647^ was administered intrathecally, and mice were decapitated, and spinal cords were extracted at different time points such as 1h, 8h, 24 h and 7 days after the tracer injection. In group 7, mice were anesthetized and focal SCI was induced, followed by an intrathecal injection of OVA^647^ 24 h later and extraction of the spinal cord 30 min after the injection of tracer. From the group 1-6, adult wildtype C57BL/6 were used whereas in group 7, C57BL/6 wildtype mice and AQP4(-/-) were used. Whole spinal cords were imaged using an Olympus epifluorescence stereomicroscope. Sections (100 µm) were cut using a vibratome and imaged. Exposure settings were kept constant for all groups. Images were acquired using either a confocal microscope (Nikon Eclipse Ti) or an epifluorescence microscope (Nikon Ni-E). Confocal acquisition settings for fluorescence were kept consistent across samples. Images were adjusted for brightness and analyzed using FIJI/ImageJ software and GraphPad Prism softwares. Approximately 10–20 sections per animal were imaged and analyzed.

### NMDA Injections and Post-Operative Care

Mice were anesthetized with a ketamine/xylazine mixture (100 mg/kg and 10 mg/kg, respectively). Lidocaine was administered locally, and buprenorphine (0.1 mg/kg, s.c.) was given as a preemptive analgesic. A midline incision was made on the back, and the lower thoracic vertebrae (T10-T12) were secured in a stereotaxic frame using spinal clamps, followed by a small laminectomy at T11 (El Waly, et al, 2021; Kaur and Berg, 2022). Control group mice (C57BL/6) received a 0.3 µL injection of phosphate-buffered saline (PBS, pH 7.4, C57-PBS) at a depth of 1 mm into the spinal cord using a glass micropipette (injection rate: 0.2 µL/min). To induce SCI in the experimental groups, which include 2 groups such as C57BL/6 wildtype mice (C57-NMDA) and AQP4(-/-) mice (AQP4KO-NMDA), 100 mM NMDA (Sigma-Aldrich Inc.) in 0.1 M PBS was injected into the spinal cord parenchyma at the same depth and rate. After injection, the needle was left in place for 10 min before being carefully withdrawn. The wound was sutured, and the animals received post-operative care. Buprenorphine mixed with Nutella (0.2 mg/1 g Nutella) was administered orally every 12 h for 3 consecutive days. Carprofen (5 mg/kg, s.c.) was given once daily for 3 days. Mice were monitored twice daily for the first 3 days followed by once daily thereafter for one week.

### Voluntary Wheel Exercise

A subset of mice was housed in polypropylene cages (36 cm L × 20 cm W × 14 cm H) equipped with a 16 cm diameter running wheel that rotated when the mouse voluntarily engaged with it. The wheel’s rotation rate was continuously recorded at 1-min intervals using magnetic switches interfaced with a PC running Vital View software (Respironics, Bend, OR). Mice in three groups: C57BL/6 mice injected with PBS (C57-PBS), C57BL/6 mice with NMDA injection (C57-NMDA), and AQP4(-/-) mice with NMDA injection (AQP4KO-NMDA) were acclimated to the exercise for one week prior to SCI induction, and they continued under the same conditions for one week post-SCI or sham surgery. Daily records were kept for total running distance, number of running bouts, and duration of the bouts, as described by Littlefield et al. (2015).

### Basso Mouse Scale (BMS)

Hindlimb locomotor function recovery post-injury was assessed using the Basso Mouse Scale (BMS) (Basso et al., 2006). Before testing, injured animals had their bladders emptied. The mice were then placed in an open field for 4 min, and locomotion scores were assessed by an observer blinded to the treatment groups.

### Quantification of Lesion Volume

To assess spinal cord damage, 100 µm-thick sections were prepared using a vibratome and stained with the Klüver–Barrera method (Luxol Fast Blue and Cresyl Violet) to detect myelin. For morphometric analysis, every third section was selected. Images were captured using an Olympus stereoscopic microscope equipped with a digital camera. Severe disruption of tissue organization and/or loss of staining was identified in the lesion area. The areas of individual sections were measured using ImageJ software, and the lesion volume was calculated by summing the total lesion areas and multiplying by the distance between sections (200 µm). The total lesion volume was defined as the sum of the lesion within the spinal cord plus any atrophic volume outside the lesion (Peng et al., 2009).

### Statistical Analysis

Data were analyzed using GraphPad Prism 7.0c. Analysed data were expressed as mean ± SEM. Statistical comparisons were made using two-way and one-way ANOVA followed by Tukey’s or Sidak’s test for multiple comparisons. Unpaired t-tests were used for all other comparisons unless otherwise indicated. A p-value < 0.05 was considered statistically significant.

## Author contributions

J.K. and M.N. developed the concept and designed the experiments. J.K., M.G., P.W. performed the experiments. J.K., M.G., D.W., D.K. analysed the experimental data. J.K. wrote the manuscript, and M.N. edited it. All authors reviewed the manuscript.

## Acknowledgements

We thank Weiguo Peng for excellent experimental work and Dan Xu for graphic support. We also thank Yuki Mori at Core Facility for Integrated BioImaging for MRI data acquisition and analysis support.

---

**Supplementary Fig. 1.**
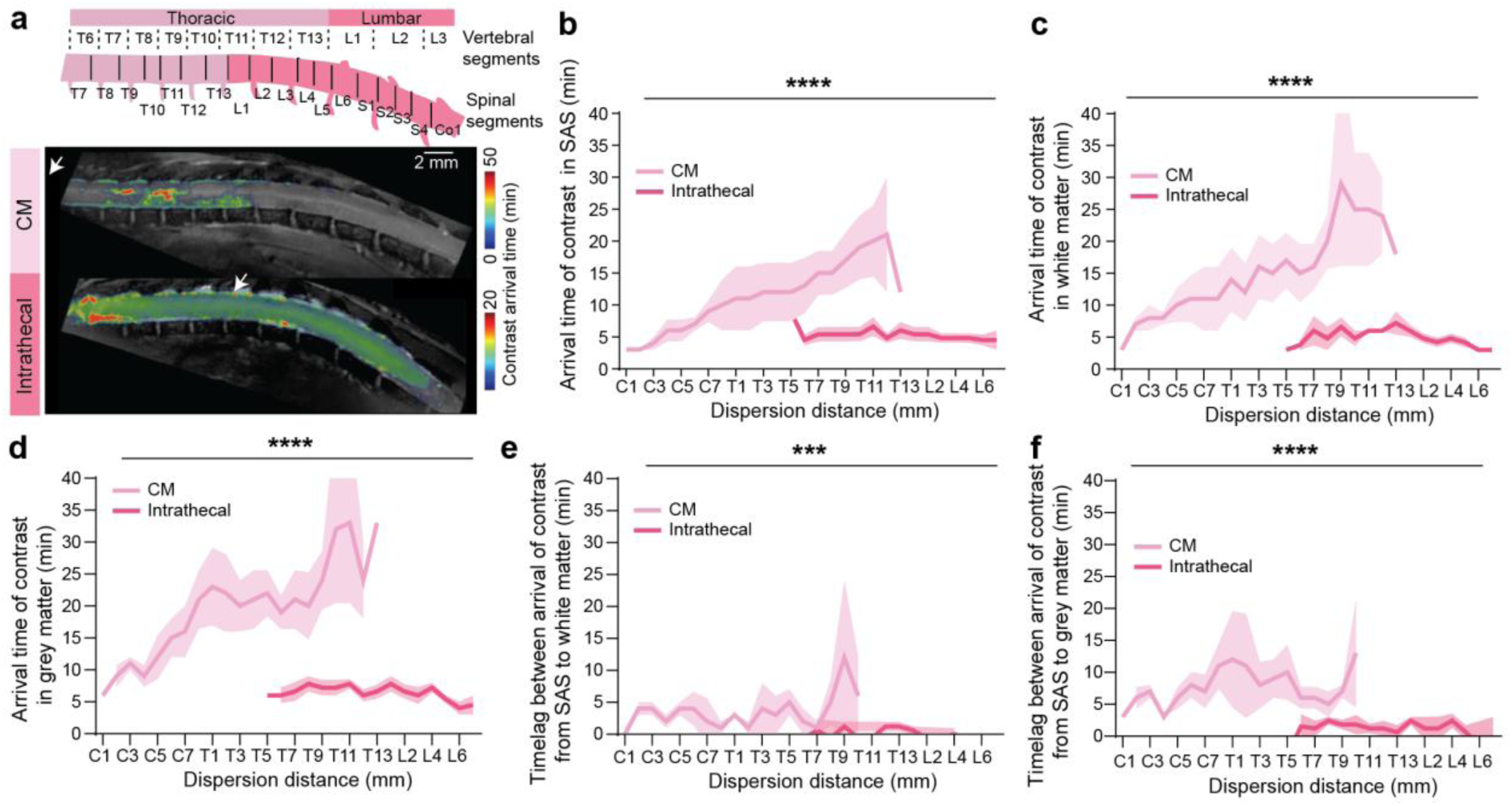
Intrathecal delivery results in faster tracer arrival compared to cisterna magna (CM) injection. **a)** Example images with segmental sketch illustrating the arrival of contrast agent in the spinal cord SAS, white and grey matter after CM or intrathecal injection. **b)** Contrast agent arrived at the C1 level of the subarachnoid space (SAS) within 3 min after the CM injection and dispersed in the various spinal levels, where it gradually arrived at the T13 level at 12 min (n=3, 2-way ANOVA, P=0.0001). Whereas after the intrathecal injection at L1 level, the contrast arrived bilaterally (at T13 level in 6.0 ± 1.2 min at L2 level in 5.4±1.0 min) and quickly dispersed in the whole thoraco-lumbar spine in an average time of 5.4±0.8 min (n=5). **c)** After CM injection, contrast arrived in white matter (WM) from SAS at C1 level within 3 min where it dispersed along the whole WM and reached at T13 level in 18 min (n=3). After intrathecal injection in L1, it dispersed bilaterally and arrived in WM T13 in 6.0±1.3 min and L2 level in 4.8±0.66 min (n = 5, 2-way ANOVA, P=0.0001). **d)** In grey matter (GM), arrival time was 6 min following CM injection and 6.6±1.1 min at T13 and 6.6±1.0 min at L2 level following intrathecal delivery at L1 spinal level (n = 5, 2-way ANOVA, P=0.0001). **e, f)** After CM injection, average time lag between the arrival of contrast in WM and GM from SAS was 3.4±0.54 min and 8.3±0.9 min, respectively. However, after intrathecal injection the average time lag for WM and GM from SAS was 0.6±0.19 min and 1.7±0.17 min, respectively (CM vs intrathecal control, 2-way ANOVA, WM: P=0.001; GM: P=0.0001).

**Supplementary Fig. 2.**
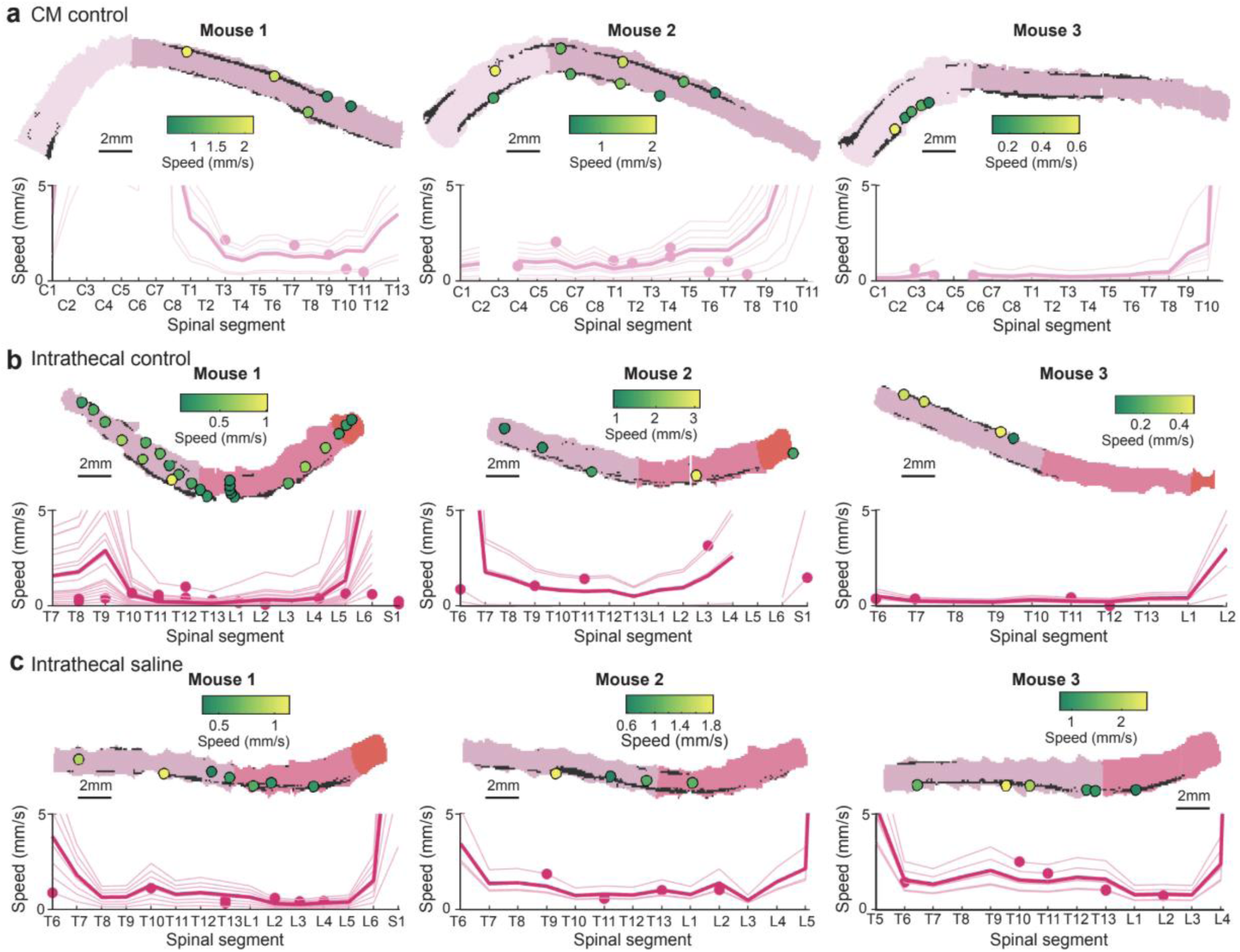
SAS volumetric analysis and speed measurements. **a) a-c)** Individual mouse data of contrast agent segment-wise speed calculation in the SAS after CM (a) or intrathecal (b-c) injection.

**Supplementary Fig. 3.**
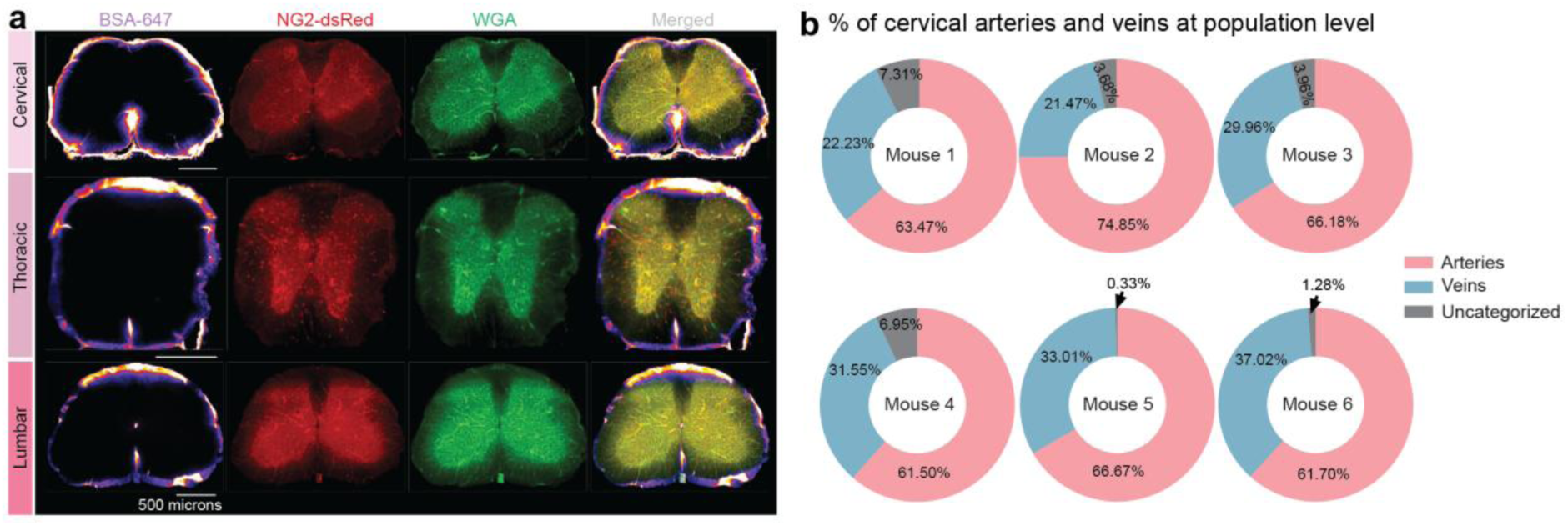
Cisterna Magna infused tracer predominantly flows along arterial perivascular spaces. **a)** Example images show spinal cord vessels labelled by bovine serum albumin conjugated with Alexa Fluor 647 (BSA^647^) and labelling of arteries and veins by wheat germ agglutinin conjugated with Alexa Fluor 488 (WGA^488^) in NG2-dsRed mice at cervical, thoracic and lumbar spinal cord levels. **b)** % of vessels in the cervical spinal cord at the population level (14-20 spinal sections per mouse) in each mouse spinal cord showed maximum % of arteries (in red) were present in the cervical region, followed by veins (in cyan) with the minimum percentage of uncategorized vessels (in grey).

**Supplementary Fig. 4.**
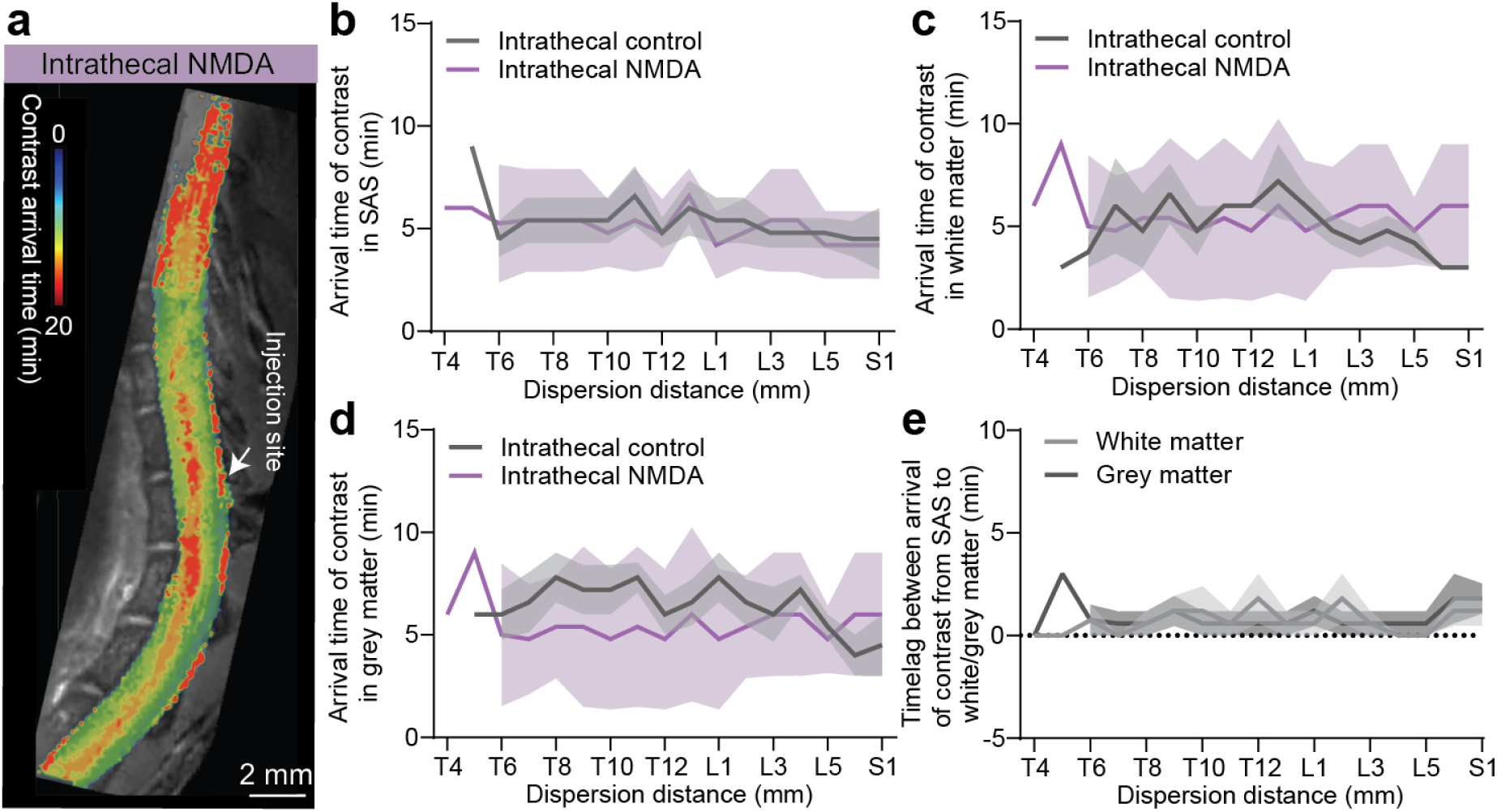
NMDA injection did not change the arrival of contrast in the spinal cord. **a)** Example image showing arrival and dispersion of contrast delivered intrathecally 24 h after NMDA injection. **b)** After intrathecal injection at L1, bilateral arrival and dispersion of contrast occurred where the contrast arrived and dispersed in the SAS quickly (rostrally at T13 within 6.0 min and caudally at L2 spinal level within 4.8±0.66 min) after SCI, which was similar to the intrathecal control conditions (n=5). **c, d)** After SCI, the contrast arrived at WM within 6.0±1.47 min at T13 level and caudally within 6.0±1.2 min at L2 level and in GM T13 level within 6.0±1.69 min and caudally at L2 level within 5.4±1.0 min (n=5). **e)** Time lag of contrast SAS to entry into the WM and GM after SCI was 0.9±0.17 min and 0.93±0.16 min, respectively (n=5). Note that no significant difference was observed in the intrathecal control (n=5) and SCI conditions in all cases.

**Supplementary Fig. 5.**
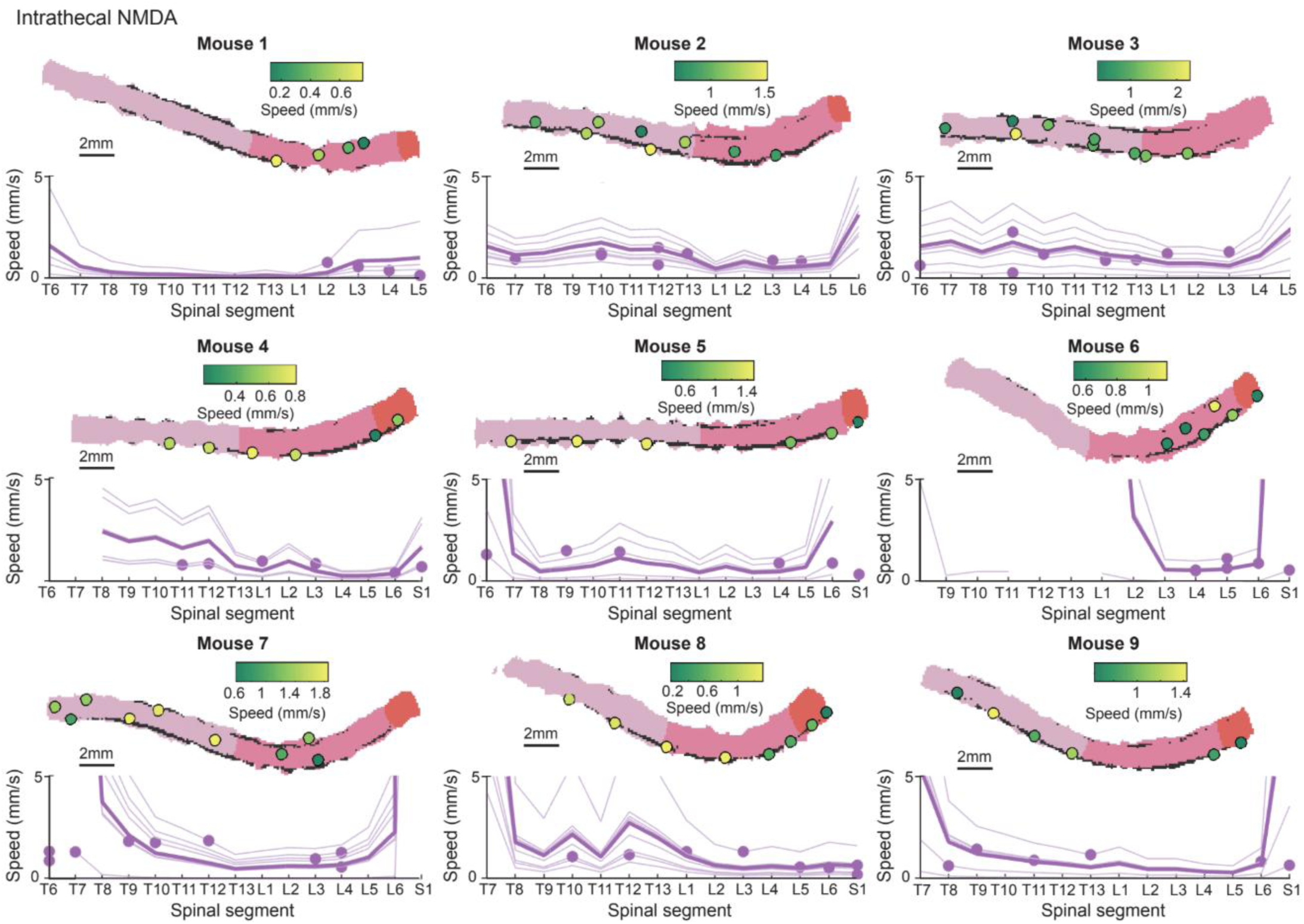
Speed measurement of contrast agent (delivered intrathecally) in the SAS 24 h after the NMDA injection. Individual mouse data from the SAS showing changes in contrast agent speed in each spinal segment after the lesion.

